# Production of Volatile Moth Sex Pheromones in Transgenic *Nicotiana benthamiana* Plants

**DOI:** 10.1101/2021.03.31.437903

**Authors:** Rubén Mateos-Fernández, Elena Moreno-Giménez, Silvia Gianoglio, Alfredo Quijano-Rubio, Jose Gavaldá-García, Lucía Estellés, Alba Rubert, José Luis Rambla, Marta Vazquez-Vilar, Estefanía Huet, Asunción Fernández-del-Carmen, Ana Espinosa-Ruiz, Mojca Juteršek, Sandra Vacas, Ismael Navarro, Vicente Navarro-Llopis, Jaime Primo-Millo, Diego Orzáez

## Abstract

Plant-based bio-production of insect sex pheromones has been proposed as an innovative strategy to increase the sustainability of pest control in agriculture. Here we describe the engineering of transgenic plants producing *(Z)*-11-hexadecen-1-ol (Z11-16OH) and *(Z)*-11-hexadecenyl acetate (Z11-16OAc), two main volatile components in many Lepidoptera sex pheromone blends. We assembled multigene DNA constructs encoding the pheromone biosynthetic pathway and stably transformed them in *Nicotiana benthamiana* plants. The constructs comprised the *Amyelois transitella AtrΔ11* desaturase gene, the *Helicoverpa armigera* farnesyl reductase *HarFAR* gene, and the *Euonymus alatus* diacylglycerol acetyltransferase *EaDAct* gene in different configurations. All the pheromone-producing plants showed dwarf phenotypes, whose severity correlated with pheromone levels. All but one of the recovered lines produced high levels of Z11-16OH but very low levels of Z11-16OAc, probably as a result of recurrent truncations at the level of the *EaDAct* gene. Only one plant line (SxPv1.2) was recovered harbouring an intact pheromone pathway and producing moderate levels of Z11-16OAc (11.8 µg g^-1^ FW), next to high levels of Z11-16OH (111.4 µg g^-1^). Z11-16OAc production was accompanied in SxPv1.2 by a partial recovery of the dwarf phenotype. SxPv1.2 was used to estimate the rates of volatile pheromone release, which resulted in 8.48 ng g^-1^ FW per day for Z11-16OH and 9.44 ng g^-1^ FW per day for Z11-16OAc. Our results suggest that pheromone release acts as a limiting factor in pheromone bio-dispenser strategies and establish a roadmap for biotechnological improvements.

## Introduction

Insect pheromones are a sustainable alternative to broad-spectrum pesticides in pest control. Different pheromone-based pest management approaches can be employed to contain herbivore populations, thus limiting damage to food, feed, industrial crops, and stored goods. These approaches include multiple strategies, such as: (i) attract-and-kill strategies, in which pheromones are used to lure insects into mass traps; (ii) push-pull strategies, in which different stimuli are used to divert herbivores from crops to alternative hosts; and (iii) mating disruption techniques in which mating is prevented or delayed by providing males with misleading pheromone cues (Cook et al., 2006). Broad-spectrum pesticides cause severe toxicity not only towards the targeted insect population but also towards predatory insects resulting in deep ecological imbalances (Witzgall et al., 2010). On the contrary, insect sex pheromones usually produced by females to attract males over long distances are highly species-specific and minimize environmental toxicity. Furthermore, pheromone-based pest control approaches are effective against pesticide-resistant insect populations and prevent the emergence of genetic pesticide resistance.

The global insect pheromone market was worth 1.9 billion USD in 2017, with projections reaching over 6 billion USD by 2025 (Agricultural Pheromones Market Report FBI100071, 2021). Despite their biological potential and their value to farmers and the environment, their use suffers from some limitations: the chemical synthesis of insect sex pheromones can often be costly and complex and generate polluting by-products, which hamper their sustainability. The cost of chemically synthesized pheromones ranges from 500 to thousands USD kg^-1^, making this solution profitable only for very high-value end products. To make pheromone production more sustainable, engineered biological systems can be designed to function as pheromone biofactories from which the molecule(s) of interest can be purified to formulate conventional traps. Ideally, live biodispensers can be envisioned, which directly release pheromones in the environment in an autonomous, self-sustained manner (Bruce et al., 2015).

Over 150,000 lepidopteran species and 1,500 lepidopteran pheromones are known (El-Sayed, 2021). Many of these butterflies and moths are relevant for agriculture and rely heavily on pheromones for mating. Lepidopteran sex pheromones have been the focus of many attempts at biotechnological production, because of their relatively simple chemical composition and their economic relevance. Sex pheromones emitted by female moths are composed of a discrete blend of volatile compounds, mostly C10-C18 straight chain primary alcohols, aldehydes, or acetates derived from palmitic and stearic acids (Löfstedt et al., 2016). Although hundreds of species share the same pheromone compounds, the components of the pheromone blend and their relative abundance constitute highly precise, species-specific cues for mating. The biosynthesis of moth pheromones is usually accomplished by three enzymatic activities. Fatty acid desaturases (FADs) introduce double bonds in specific positions (the most common in Lepidoptera are Δ9 and Δ11). Fatty acyl reductases (FARs) produce fatty alcohols, with different substrate specificities (some accept only a limited range of substrates, while others are more promiscuous). Finally, most moth pheromones are obtained through esterification of fatty alcohols, to produce aldehydes and acetates. Acetyltransferases are thought to perform this latter step, although no insect acetyltransferases have been identified which work on fatty alcohols (Petkevicius et al., 2020).

Plants represent an alluring platform to produce moth sex pheromones: the scalability and relatively low costs and infrastructure requirements of plant biofactories make this system versatile and sustainable. In plants, fatty acid biosynthesis starts in the chloroplast relying on the availability of C3 products generated by photosynthesis. Fatty acids are then modified (by desaturation, chain length modification, isomerization or substitution), and precursors are exported to the cytosol, where the final biosynthetic steps take place. In a pioneer work, Nešněrová et al., (2004), took advantage for the first time of the plant fatty acids pool to produce lepidopteran pheromones precursors in plants. Later, in the most extensive screening of candidate genes so far, Ding et al., (2014) identified the most effective among 50 different gene combinations to produce moth pheromones by transient expression in *N. benthamiana*. Subsequently, Xia et al., (2020) established stably transformed *N. benthamiana* and tobacco lines to express precursors for the synthesis of a wide range of moth pheromones. However, to this date, no stable transgenic plants have been reported producing the actual volatile pheromone components.

In this work, we aimed to test the ability of *N. benthamiana* plants to act as constitutive moth pheromone biofactories. *Nicotiana* species (*N. tabacum* and *N. benthamiana*) are ideal chassis for metabolic engineering, due to their large leaf biomass (especially for plastid-derived products) and amenability to genetic manipulation, both through stable transformation and agroinfiltration. For stable pheromone production we selected three of the genes identified by Ding et al., (2014), namely the *Amyelois transitella AtrΔ11* reductase, the *Helicoverpa armigera* reductase *HarFAR* and the plant diacylglycerol acetyltransferase *EaDAct* from the bush *Euonymus alatus*. The products of this pathway, *(Z)*-11-hexadecen-1-ol (Z11-16OH) and its ester *(Z)*-11-hexadecenyl acetate (Z11-16OAc), are components of the specific pheromone blends of over 200 lepidopteran species (El-Sayed, 2021). The generation of transgenic pheromone-producing plants (named SxP plants) turned out to be severely hampered by a strong growth penalty putatively imposed by the pheromone biosynthetic pathway. In a first round of attempts, only *N. benthamiana* plants accumulating the fatty alcohol were recovered, with all primary transformants showing dwarf phenotypes in different degrees. New strategies aimed at ensuring the integrity of the construct finally yielded a single transgenic line accumulating both the alcohol and the acetate components at relatively high levels, while maintaining acceptable levels of fertility and biomass production. This single line allowed us to gain insights into the challenges associated with fatty-acid derived pheromone production in plants, such as yield-associated growth penalties, changes in volatile profile, and compound volatility.

## Results

### Assembly of the metabolic pathway

To assess plant-based production of the target moth sex pheromones (Z11-16OH and Z11-16OAc), a T-DNA construct encoding the three biosynthetic genes under the control of constitutive CaMV35S promoters, plus a silencing suppressor (P19), was agroinfiltrated in plant leaves (Figure 1(a)), and the total volatile organic compound (VOC) composition was analysed at 5 days post infiltration by gas chromatography/mass spectrometry (GC/MS). GC peaks corresponding to the pheromone compounds Z11-16OH and Z11-16OAc were detected in samples transformed with all three enzymes, but not with P19 alone (Fig. 1(b)). Moreover, both substances were among the most predominant compounds in the leaf volatile profile, indicating that the transgenes were expressed at high levels. Interestingly, a small peak identified as *(Z)*-11-hexadecenal (Z11-16Ald) was also detected in the agroinfiltrated samples, likely due to the endogenous activity of alcohol oxidases, as previously suggested by Hagström et al., (2013). This aldehyde is itself a component of the pheromone blends of around 200 lepidopteran species (El-Sayed, 2021). Based on these results, a multigene construct for stable transformation of *N. benthamiana* plants was assembled (GB1491). This construct comprised the three constitutively expressed enzymes, the kanamycin resistance gene *NptII* and the visual selection marker *DsRed* (Fig. 1(c)). Plants resulting from this transformation were identified as the first version of the pheromone-producing plant (SxPv1.0).

**Figure 1.**
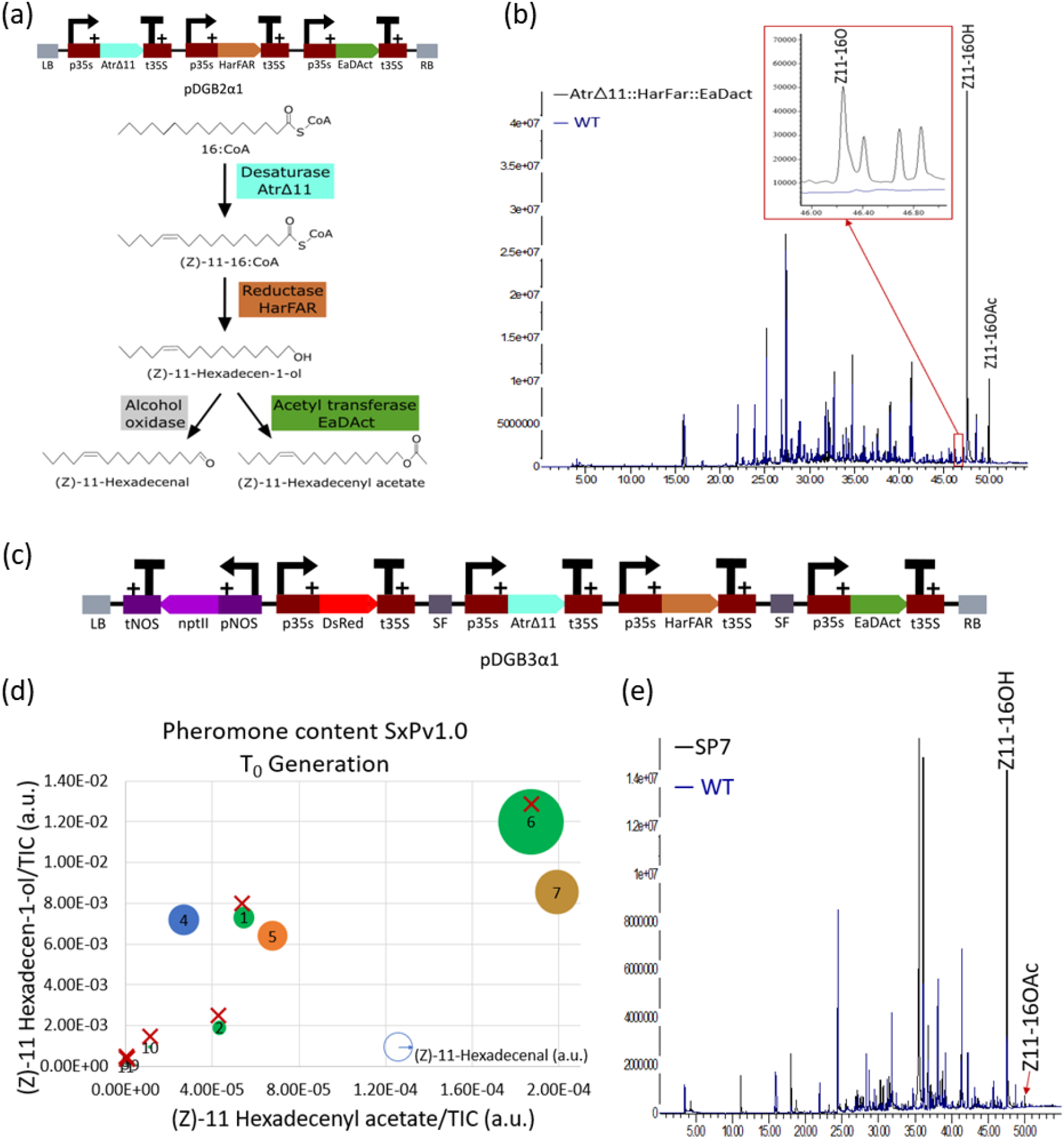
Stable and Transient expression in *Nicotiana benthamiana* of the synthetic moth pheromone pathway. (a) Schematic view of the T-DNA construct used for transient expression, carrying the three transgenes *AtrΔ11, HarFAR* and *EaDAct*, each under the control of the constitutive CaMV35s promoter and terminator, and the biosynthetic route of the moth pheromones. (b) GC/MS analysis of the volatile profile of *N. benthamiana* transiently expressing the transgenes (black line) and a mock infiltrated plant with only P19 (blue line). Peaks corresponding to the target insect pheromones are indicated with a label. Highlighted in red is the region of the (*Z*)-11-hexadecenal peak. (c) Schematic view of the T-DNA construct for the SxPv1.0 encoding the three transgenes and two selection markers; (d) Pheromone content in SxPv1.0 T_0_ plants (numbered from 1 to 11). The width of each dot corresponds to the (*Z*)-11-hexadecenal level of each sample. Plants marked with a red cross died before seeds could be collected. (e) Overlapped chromatograms showing the volatile profile of a representative SxPv1.0 T_0_ plant (black line) and a WT plant (blue line).

### SxPv1.0 stable transformants

The transformation of *N. benthamiana* with the GB1491 construct resulted in the selection of 11 kanamycin-resistant shoots, which also showed *DsRed* fluorescence (T_0_ generation SxPv1.0 plants). These shoots were further grown and rooted, and leaf samples were collected at the early flowering stage to assess pheromone production. As observed in Figure 1(d), several T_0_ plants presented detectable levels of all three pheromone compounds in variable amounts. A correlation was readily observable in the relative abundance of all three pheromone compounds in each plant, despite Z11-16OAc levels being much lower than expected in all cases compared to transient pheromone expression. Furthermore, although phenotypic evaluation of *N. benthamiana* T_0_ lines is generally cumbersome due to the influence of *in vitro* culture, severe growth penalties were observed in these plants, and only 5 out of 11 plants (SxPv1.0_4, 5, 7, 8, and 9) survived long enough to produce seeds.

To further understand the phenotypic effects of pheromone production, the progeny of plants SxPv1.0_4, SxPv1.0_5 and SxPv1.0_7 was analysed up to the T_3_ generation and the plant size and pheromone production levels were recorded for each individual. In the T_1_ generation, growth penalties were also observed in most descendants for all three lines, generally associated with high pheromone production (Fig. 2(a) and (b), left plot). Several plants could not be phenotyped as they died soon after germination. Those producing enough seeds were brought to T_2_, where a similar trend was also observed (Fig. 2(b), central panel). A few T_2_ plants clearly separated from the rest in terms of high Z11-16OH production, which was again associated with small size and reduced fertility. Neither T_1_ or T_2_ plants showed signs of recovery in Z11-16OAc levels, although the corresponding GC/MS peak remained detectable and above the wild type (WT) baseline (not shown). At this stage, we decided to re-evaluate the integrity of the T-DNA in T_2_ plants, finding that DNA rearrangements had occurred in all three lines at the *EaDAct* site, resulting in a truncated gene. Interestingly, at least two independent truncation events could be inferred from PCR analysis of gDNA and cDNA samples. In SxPv1.0_7_4 plants, the presence of a ∼700bp insertion of a DNA fragment of plasmid origin could be determined at the 3′ end of the *EaDAct* coding sequence. In contrast, the same 700bp genomic PCR fragment could not be recovered from the offspring of SxPv1.0_4_2, SxPv1.0_5_1 and SxPv1.0_5_2 plants, which nevertheless had also a truncated ORF as evidenced by PCR analysis of cDNA samples (Supplementary Fig. 2).

**Figure 2.**
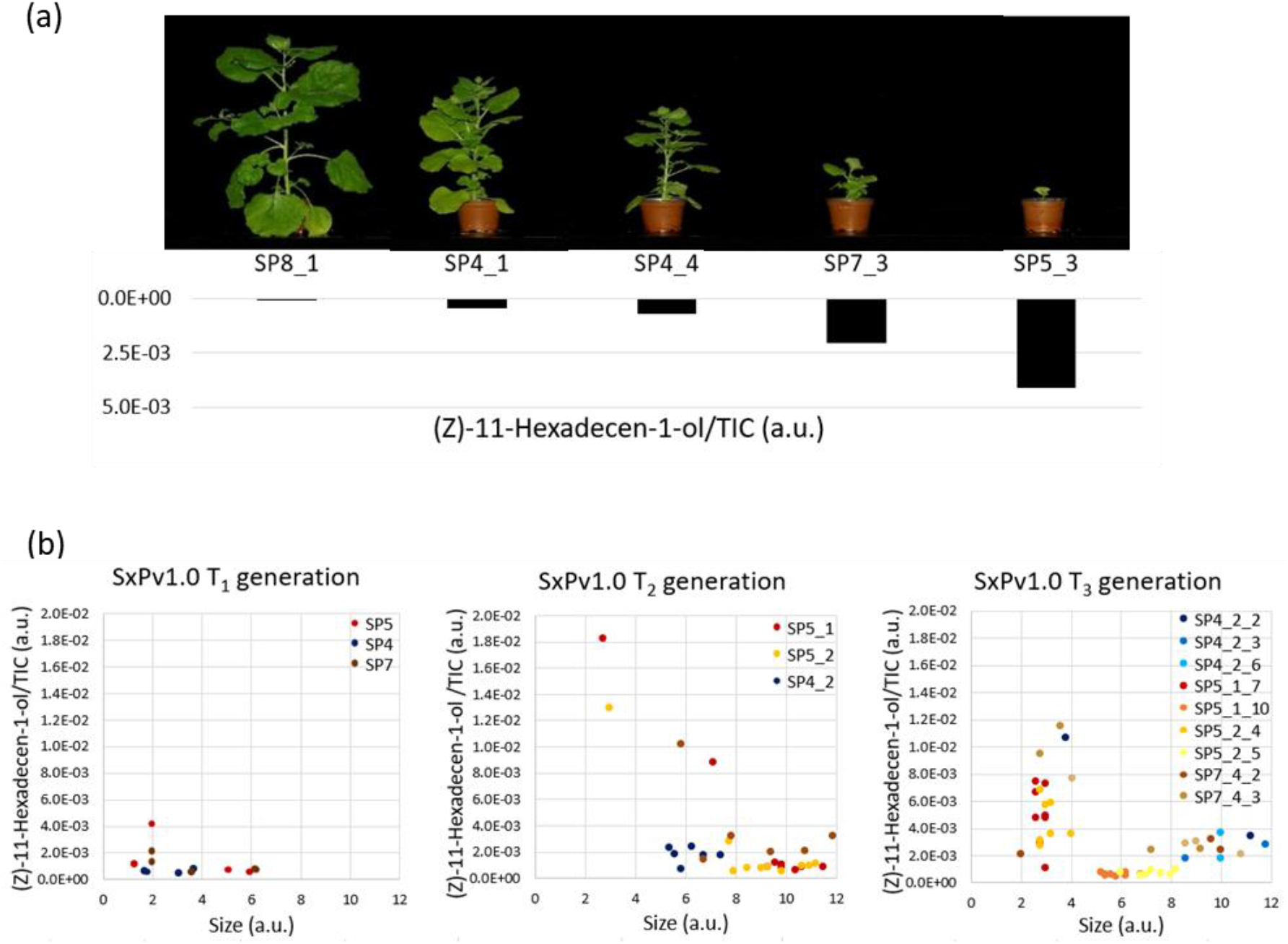
Growth penalty linked to pheromone production in SxPv1.0 plants. (a) Representative SxPv1.0 T_1_ plants with their corresponding (*Z*)-11-hexadecen-1-ol levels. (b) Correlation between (*Z*)-11-hexadecen-1-ol content (arbitrary units, a.u.) and plant size (a.u.) in all three generations of SxPv1.0.

Despite the detection of a T-DNA truncation, the analysis of SxPv1.0 offspring was continued up to T_3_ (Fig. 2(b), right plot), where a sharp separation between low and high producers was consolidated. Interestingly, the offspring from the SxPv1.0_5_1_7 homozygous line (100% kanamycin resistant) comprised only high producer plants, whereas heterozygous lines as SxPv1.0_4_2_2 or SxPv1.0_7_4_3 segregated in high and low producers, these correlating with small and large sized individuals, respectively. This observation strongly indicates a drastic effect of transgene copy number in both growth and pheromone production.

### New stable transgenic versions SxPv1.1 and SxPv1.2

The presence of at least two independent truncation events affecting *EaDAct* prompted us to design new transformation strategies by placing a selection marker adjacent to the *EaDAct* gene, ensuring its integrity. Two new DNA constructs were assembled (SxPv1.1 and SxPv1.2) carrying *DsRed* and *nptII* at different relative positions of the T-DNA, as depicted in Fig. 3(a), (b). Five SxPv1.1 and eight SxPv1.2 kanamycin resistant T_0_ plants were recovered for each transformation, many of them showing detectable red fluorescence, but unfortunately all but one failed to produce detectable levels of pheromones. The only exception corresponded to plant SxPv1.2_4, which showed Z11-16OH and Z11-16Ald amounts comparable to the SxPv1.0 plants, but also Z11-16OAc levels close to those measured in transient experiments (Fig. 3(c), (d)). SxPv1.2_4 presented premature flowering, a feature that is not unusual in T_0_ *N. benthamiana* plants, and produced viable seeds, giving us the opportunity to further investigate the phenotype of stable Z11-16OAc producers.

**Figure 3.**
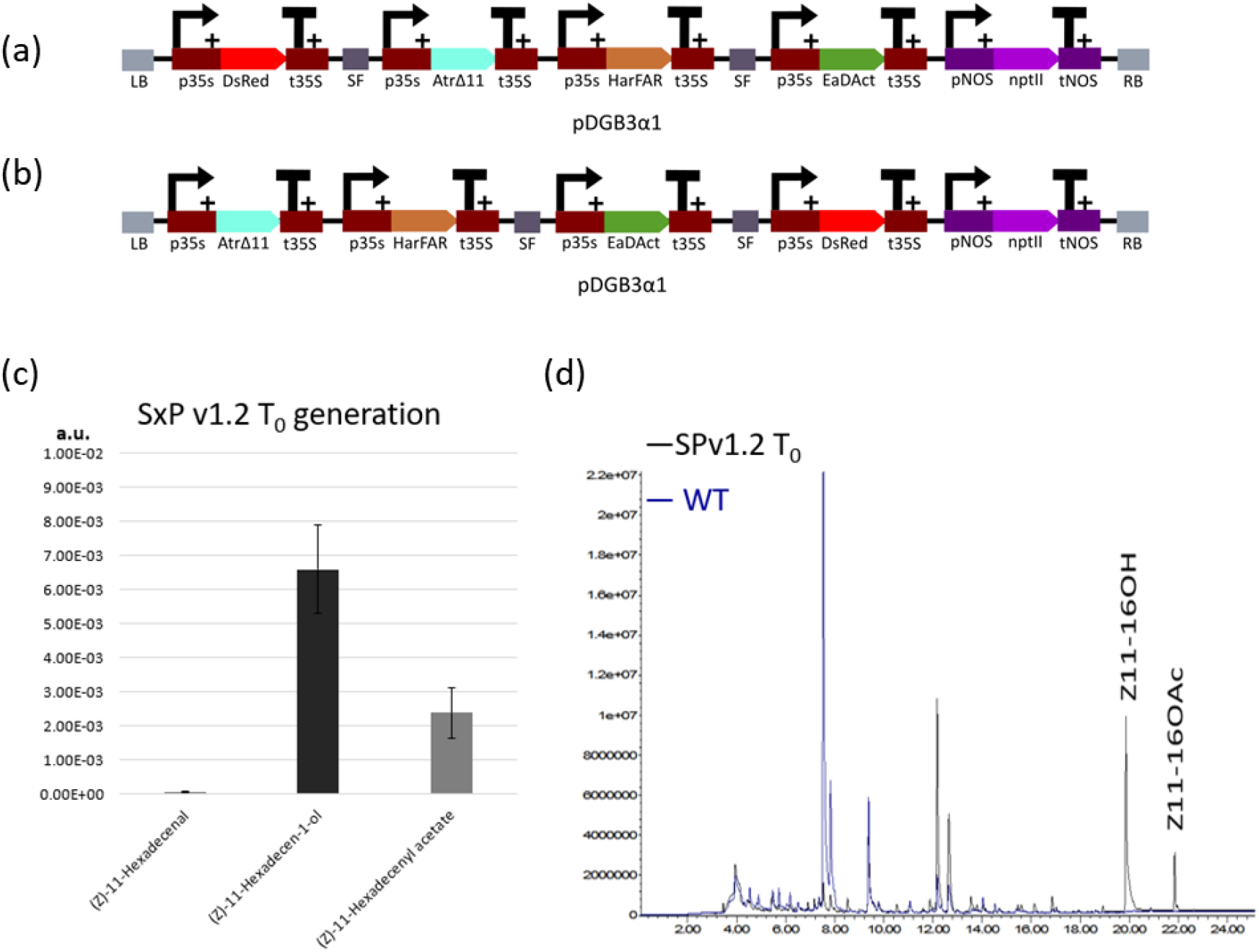
SxP version 1.1 and 1.2 stable plants. (a) Schematic representation of the T-DNA construct employed for stable transgenic SxPv1.1; (b) Schematic representation of the T-DNA construct employed for stable transgenic v1.2; (c) Pheromone content in the surviving SxPv1.2 T_0_ plant. Error bars represent the average ± SE of 3 independent replicates; (d) Overlapped chromatograms showing the volatile profile of a SxPv1.2 T_0_ plant (black line) and a WT *Nicotiana benthamiana* (blue line).

For a deeper understanding of the effect of fatty-acid-derived pheromone production in plant homeostasis, a comparative study between the progeny of the T_2_ SxPv1.0_5_1_7 homozygous line and the T_0_ SxPv1.2_4 line was performed. All analyzed SxPv1.2 T_1_ seeds (>50) were kanamycin resistant, indicating multiple copy insertions. The relative levels of all three pheromone compounds in leaves at two different developmental stages (juvenile and adult) was recorded for twelve T_1_ plants per genotype. Similarly, pheromone content in roots was also measured at the adult stage. Plant size was recorded for all analyzed individuals. As expected, all transgenic plants produced detectable levels of both pheromones but only in the case of SxPv1.2, Z11-16OAc and Z11-16OH accumulated at similar levels. In all SxPv1.2 samples, the higher Z11-16OAc accumulation seems to result in lower precursor alcohol levels, compared with equivalent SxPv1.0 samples. Both insect pheromones are produced at higher levels in adult plant leaves (Fig. 4(b)) when compared with young plant leaves (Fig. 4(a)) and roots (Fig. 4(c)). All pheromone-producing plants showed considerably reduced plant size, however the growth penalty was significantly more pronounced in plants accumulating mainly Z11-16OH, whereas the conversion into the acetate form in SxPv1.2 seems to partially relieve the dwarf phenotype. Interestingly, both SxPv1.0 and SxPv1.2 plants showed similar morphology, with short petioles curved upwards and resulting in a compact “cabbage-like” characteristic shape (Fig. 4(d)). Both SxP lines showed early senescence symptoms, with premature and progressive yellowing, which led, in the case of SxPv1.0, to the premature death of the plant soon after fruit set.

**Figure 4.**
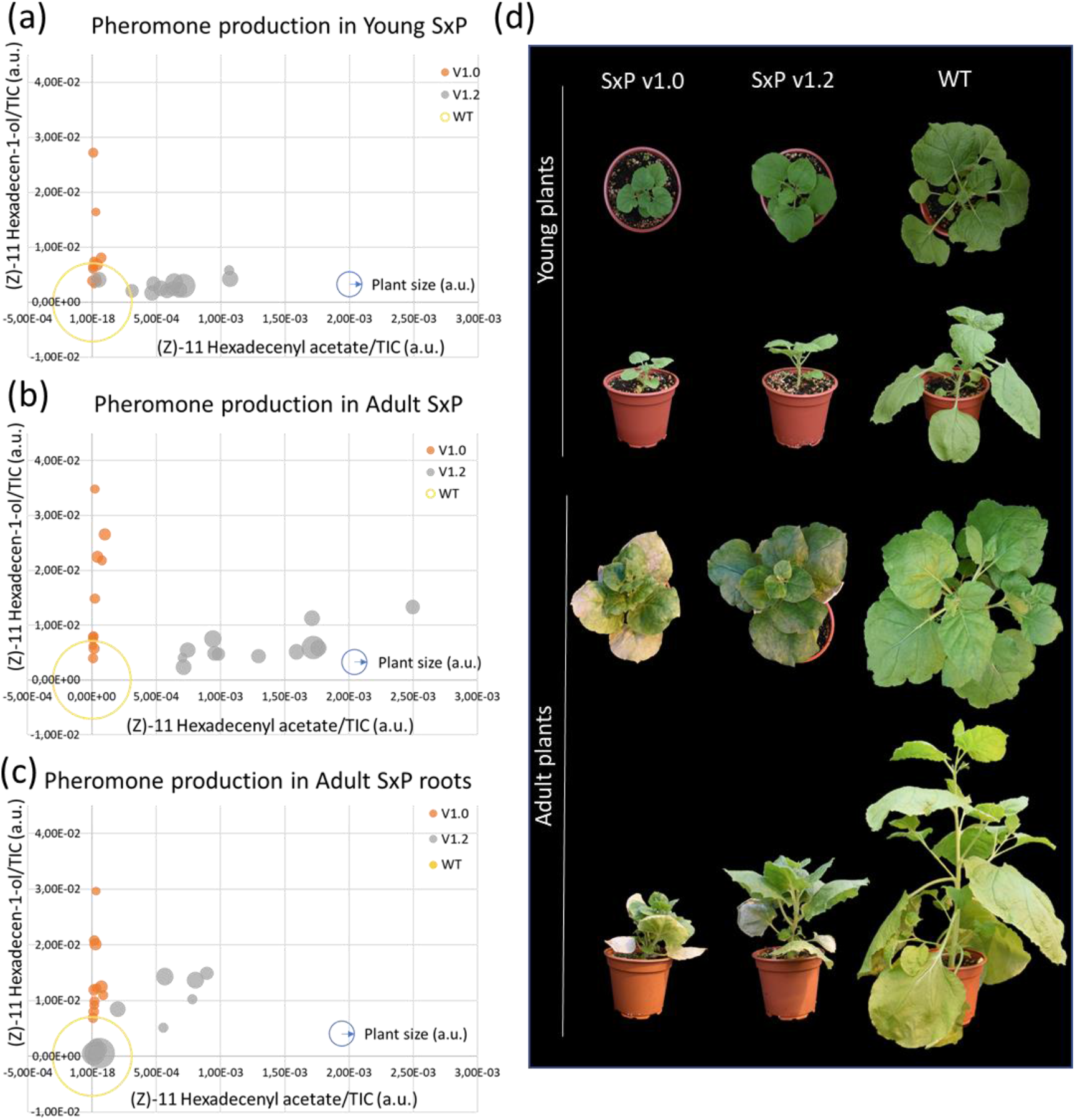
Comparative study between SxPv1.0 T_3_ and SxPv1.2 T_1_ plants. (a) Pheromone content in leaf samples from young plants of WT, SxPv1.0 and SxPv1.2 lines; (b) pheromone content in leaf samples from adult plants of WT, SxPv1.0 and SxPv1.2 lines; (c) pheromone content in root samples from adult plants of WT, SxPv1.0 and SxPv1.2 lines. The radius of each dot corresponds to the plant size of each sample. Empty circles correspond to WT plants. (d) Comparative physiological development between SxPv1.0 5_1_7_X (T_3_), SxPv1.2 4_X (T_1_) and WT *Nicotiana benthamiana* plants at the young and adult stage. Pictures were taken from representative individuals at juvenile (4 weeks after transplant) and adult (7 weeks after transplant) stages.

### The plant volatilome is affected by pheromone production

A non-targeted analysis of the plant volatile profiles was undertaken to understand the influence of the engineered pheromone pathway on the volatilome. The analysis included leaf samples of 12 young and 12 adult plants from the progeny of SxPv1.0 5_1_7, SxPv1.2_4 and the wild type. The Principal Component Analysis score plot based on the volatile profile showed clustering of the samples based on the different sample classes (Fig. 5(a)). The first component accounted mainly for differences in leaf age, whereas the second principal component separated samples according to their genotype. Remarkably, SxPv1.2 samples have, according to both components, intermediate characteristics between the WT and SxPv1.0. The greater separation of SxPv1.0 and the WT probably reflects the more deleterious phenotypic effects experienced by lines accumulating higher Z11-16OH levels.

**Figure 5.**
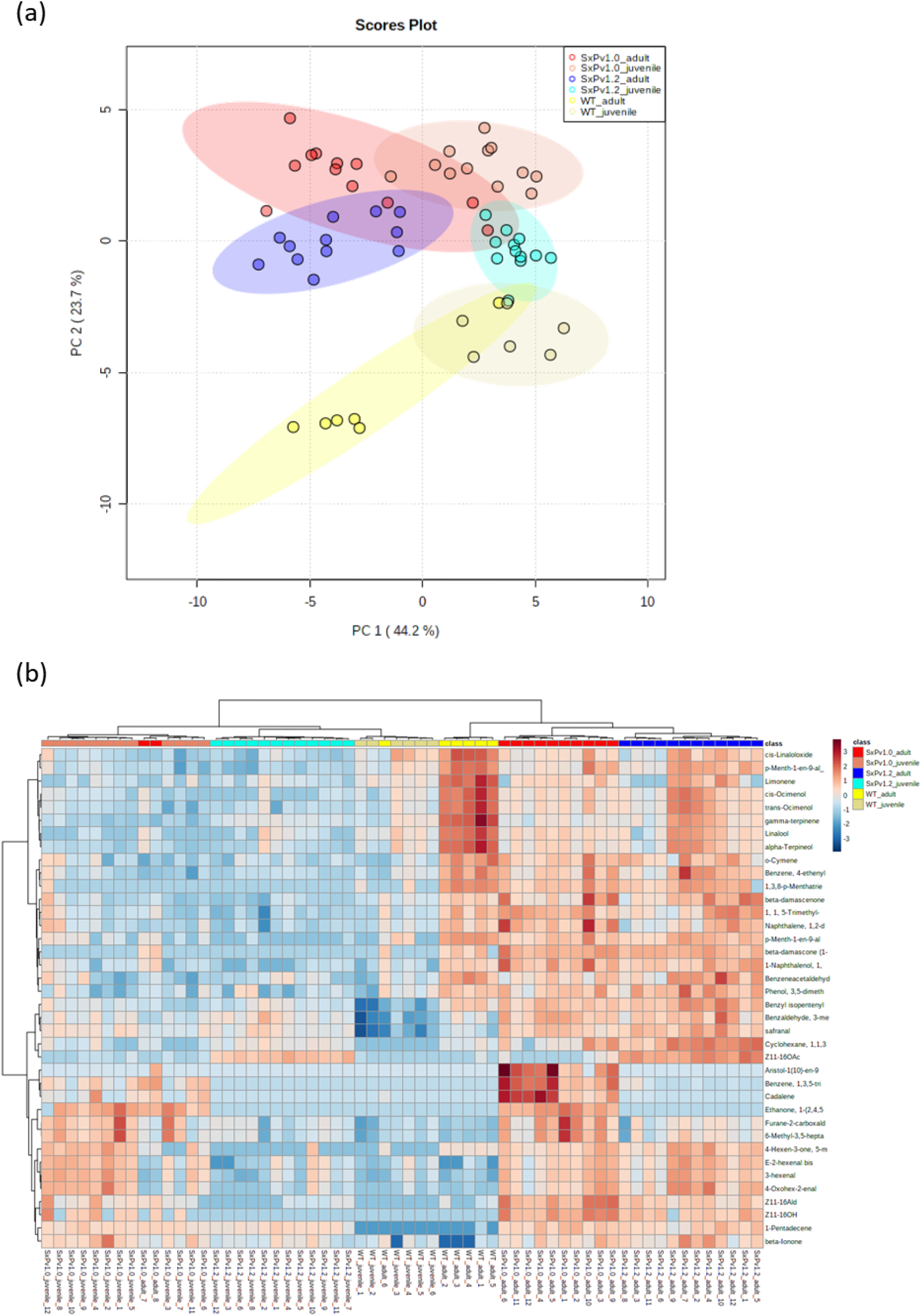
Untargeted analysis of the volatilome of SxPv1.0 and SxPv1.2, and of WT *N. benthamiana*. (a) Principal Component Analysis and (b) Hierarchical clustering and heatmap representation (obtained using Ward’s minimum variance method and Euclidean distance) of the composition of the volatilome of SxPv1.0, SxPv1.2 and wild type *N. benthamiana* leaves. Twelve individuals for each SxP genotype and six WT plants were analyzed at two developmental stages, young (4 weeks old) and adult (7 weeks old).

A clustered heatmap provides interesting visual information on the volatile leaf profiles of SxP plants (see Fig. 5(b)). The clustering reproduces with few exceptions the different classes, indicating that each genotype and each developmental stage produces a differential and characteristic blend of VOCs. In addition to the pheromone compounds themselves, which are clearly clustered in their respective groups, all SxP plants differentially accumulate other fatty-acid-derived volatile compounds (e.g. (E)-2-hexenal, 1-pentadecene), indicating a general activation of this metabolic pathway. Some compounds are characteristic of the adult stage, independently of genotype. This is the case for some apocarotenoids such as β-damascenone and β-damascone, and some phenylalanine-derived compounds such as o-cymene and phenylacetaldehyde. Other VOCs, such as monoterpenoids (α-terpineol, linalool, limonene and ocimenol), are markedly more abundant in WT than in SxP leaf tissues, with a gradient in which SxPv1.2 shows intermediate features between the wild type and the SxPv1.0 genotype. On the other hand, SxPv1.0 plants display a specific subset of volatile compounds (including the sesquiterpene cadalene) which accumulate at increasing levels at the adult stage. Z11-16Ald is detectable in both SxPv1.0 and SxPv1.2, although its levels are higher especially in leaves from adult SxPv1.0 plants, correlating with higher Z11-16OH production. In SxPv1.2 plants, in which Z11-16OH is partially converted to Z11-16OAc, Z11-16Ald is present at lower levels. Z11-16OAc is, instead, clearly restricted to SxPv1.2. The levels of all three pheromones increase with plant age.

### Pheromone identification and determination of its biological activity

Samples of Z11-16OH, Z11-16OAc and Z11-16OAld were synthesized and characterized by GC/MS and nuclear magnetic resonance (NMR) to have analytical standards of the biosynthetic targets. Additionally, to provide unequivocal identification of the plant-made compound, hexane extracts of 120 g of SxPv1.2 leaves were purified by gravity column chromatography after solvent evaporation, and a 2-mg sample of the purest fractions of the biosynthesized alcohol (Z11-16OH) was also analysed using NMR. The purity assigned by GC/MS was ca. 78 % (Supplementary Figure 3), and the data extracted from the main signals of both ^1^H and ^13^C NMR spectra were fully consistent with those obtained for the synthetic sample of Z11-16OH, confirming the structure and the cis-configuration of the double bond (Supplementary Figures 4-5). Further confirmation of the biological activity was provided by electrophysiological analysis. Hexane extracts of SxPv1.2 leaves were fractionated by column chromatography and the fractions were analysed by GC/MS. Those fractions that mainly contained Z11-16OH were gathered and employed in electroantennography (EAG) assays with *Sesamia nonagrioides* male moths. The plant-made pheromone was active, since the EAG probe registered significant antennal depolarizations when Z11-16OH reached the antennal preparations (Fig. 6; retention time 15.89 min).

**Figure 6.**
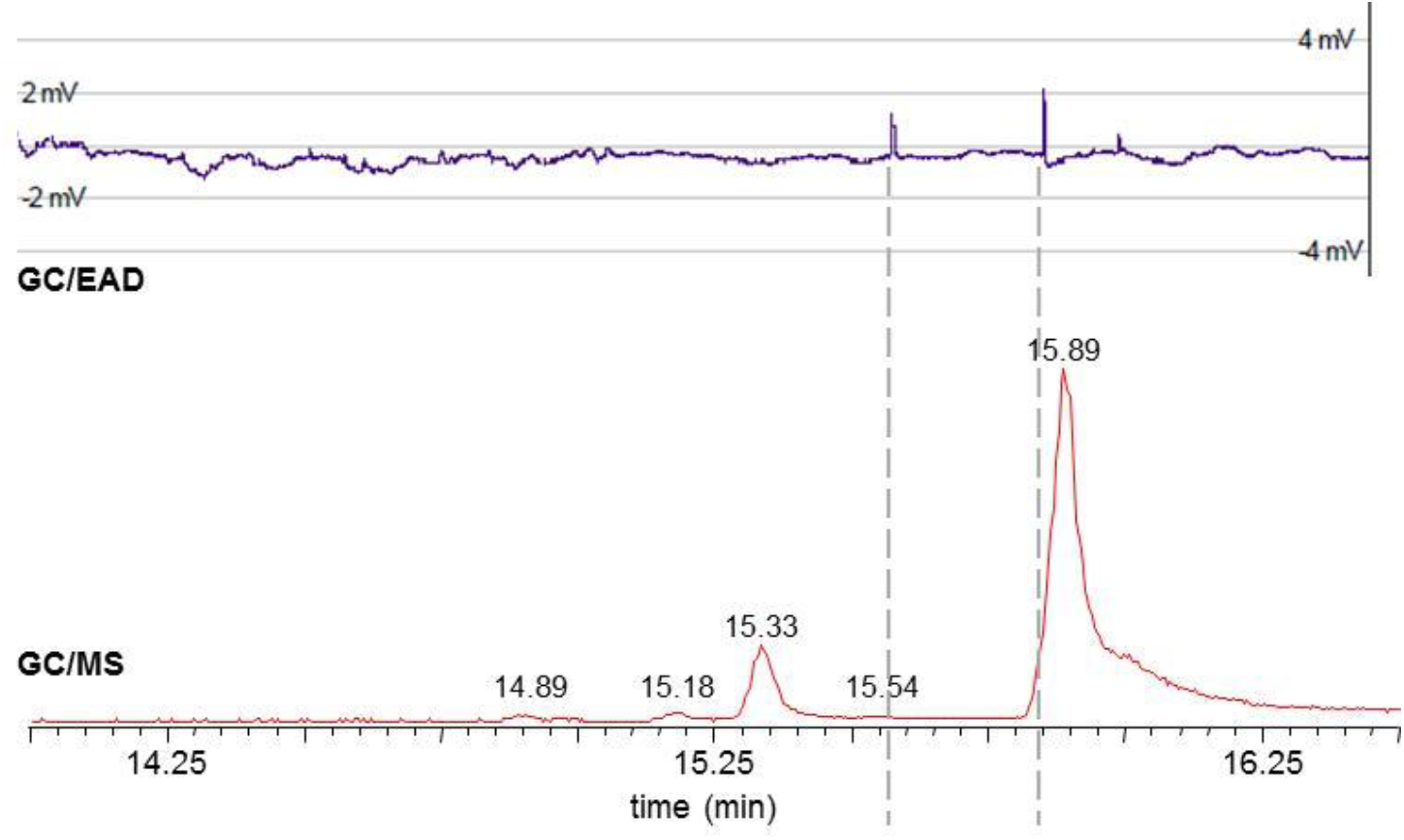
Electrophysiological activity. GC/MS chromatogram and EAD recording showing *Sesamia nonagrioides* male antenna response to the biosynthetic Z11-16OH (retention time = 15.89 min). Other compounds contained in the tested fraction were able to interact with the antennal receptors and triggered antennal responses (e.g. retention time 15.54).

### Quantification of total pheromone contents and release

The total pheromone content and the rate of volatile emission were both quantified in SxPv1.2 plants. Solvent extraction was carried out in fresh leaves as well as in leaves stored at -20°C and -80°C to evaluate the total content and the possible loss of pheromone under different storage conditions (Table 1). The Z11-16OH content in leaves was found to be in the range of 0.1 mg g^- 1^, (average 111.4 ± 13.7 µg g^-1^), whereas Z11-16OAc accumulated at lower levels (average 11.8 ± 1.3 µg g^-1^). Both pheromones were preserved in frozen leaves, although a part of Z11-16OH could be lost upon storage. Interestingly, the unbalanced ratio between the two main compounds was compensated when the rate at which pheromones are released to the environment was estimated. As shown in Table 2, an adult SxPv1.2 plant releases on average 79.3 ± 6.3 ng of Z11-16OH and 88.3 ± 11.5 ng of Z11-16OAc per day, as estimated in volatile collection experiments carried out in dynamic conditions. Not unexpectedly, this indicates a much higher volatility of the acetylated moiety. In terms of pheromone release per biomass unit, both compounds are emitted at levels close to 0.01 µg day^-1^ g^-1^ (FW). Remarkably, the emitted pheromones represented roughly 0.01-0.1% (Z11-16OH and Z11-16OAc, respectively) of the total amounts produced and accumulated in the leaves.

**Table 1.** **Quantity (µg) of Z-11-hexadecen-1-ol (OH) and (Z)-11-hexadecenyl acetate (OAc) extracted from SxPv1.2 individuals by solvent extraction and GC/MS/MS quantification.**

| plant | material | $\mu\text{g OH/g plant}$ | $\mu\text{g OAc/g plant}$ |
| --- | --- | --- | --- |
| SxPv1.2 T<sub>1</sub>-3 | fresh leaves | 164,9 | 9,6 |
| SxPv1.2 T<sub>1</sub>-4 | fresh leaves | 129,9 | 8,6 |
| SxPv1.2 T<sub>1</sub>-5 | frozen -20C | 78,1 | 10,5 |
| SxPv1.2 T<sub>1</sub>-6 | frozen -20C | 115,1 | 17,3 |
| SxPv1.2 T<sub>1</sub>-7 | frozen -80C | 75,8 | 11,8 |
| SxPv1.2 T<sub>1</sub>-8 | frozen -80C | 104,8 | 12,9 |
| | mean ± se | 111.4 ± 13.7 | 11.8 ± 1.3 |

**Table 2.** **Quantity (ng) of Z-11-hexadecen-1-ol (OH) and (Z)-11-hexadecenyl acetate (OAc) released by SxPv1.2 individuals obtained by volatile collection and GC/MS/MS quantification.**

| plant | ng collected OH | ng OH/day | ng collected OAc | ng OAc/day |
| --- | --- | --- | --- | --- |
| SxPv1.2 T<sub>1</sub>-1 | 209,3 | 69,8 | 193,3 | 64,4 |
| SxPv1.2 T<sub>1</sub>-2 | 225,6 | 75,2 | 359,4 | 119,8 |
| SxPv1.2 T<sub>1</sub>-3 | 222,6 | 74,2 | 250,7 | 83,6 |
| SxPv1.2 T<sub>1</sub>-4 | 293,9 | 98,0 | 256,8 | 85,6 |
| mean ± se | 237.8 ± 19.0 | 79.3 ± 6.3 | 265.0 ± 34.6 | 88.3 ± 11.5 |

## Discussion

This research was initiated as a Synthetic Biology project in the frame of the iGEM competition, where undergraduate students proposed the use of genetically engineered plants as dispensers of insect sex pheromones. The manufacturing of pheromones and their precursors employing biological factories such as microbial bioreactors (Hagström et al., 2013; Holkenbrink et al., 2020) or plant biofactories (Ding et al., 2014; Ortiz et al., 2020; Xia et al., 2020) has become an intensively pursued objective, fuelled by the expected gains in sustainability. Beyond the general biofactory concept, our envisioned long-term approach consists in the design of plants that function as autonomous bio-dispensers of semiochemicals. A remarkable precedent of this concept was the engineering of wheat plants releasing the alarm pheromone *(E)*-β-farnesene as a protective strategy against aphid infestation (Bruce et al., 2015). Differently to the alarm pheromone concept, which was produced in the crop itself, the proposed bio-dispenser (originally named as “Sexy Plant”, SxP) is based on the intercropping strategy where a companion crop, rather than the main crop, is engineered to emit the sex pheromone into the environment. From here, two different strategies can be followed. In a mating disruption strategy (Carde and Minks, A K, 1995; Stelinski et al., 2013; Benelli et al., 2019), the dispensers release pheromones at relatively large quantities, impairing males’ ability to detect females and therefore disrupting the mating. Oppositely, in mass trapping or attract-and-kill strategies (Hossain et al., 2006), dispensers release pheromones to attract males to traps. This later approach often requires lower pheromone levels to be released in the environment, but in turn requires higher semiochemical specificity (in terms of isomeric purity and exact ratios of the pheromone components), and also some associated equipment to trap and eventually kill the attracted insects.

The genetic engineering of *N. benthamiana* shown here was inspired by the seminal work by Ding et al., (2014), where transient expression of various components of moth sex pheromone blends was achieved. Contrary to other insect pests whose sex pheromones are made of a single, highly specific molecule, as with some mealybugs (Zou and Millar, 2015), lepidopteran sex pheromones are often made of more complex blends of fatty-acid derived compounds, many of them shared by several species. Species-specificity in these cases is provided by the precise ratio in the blend. For instance, *Sesamia nonagrioides* (Lefèbvre) males are attracted to a Z11-16Ald:Z11-16OAc:Z11-16OH blend in a 77:8:10 proportion (Sans et al., 1997). This feature makes the genetic design of plant emitters for attract-and-kill strategies in moths extremely challenging, because ensuring the right proportions of the three compounds requires a tight control of several factors, from gene expression to enzymatic activity and differential release ratios. Conversely, mating disruption seems a more attainable objective in terms of pheromone heterologous production since, in many cases, the release of non-attractive incomplete mixtures can disrupt mating as effectively as the complete blend (Evenden, M., 2016). In this case however, the main requirement imposed on a biological dispenser is to produce and release enough quantities of one or more compounds in the blend. Therefore, the main objective of this work was to understand the biological constraints accompanying the production and release in *N. benthamiana* plants of two of the most representative compounds in lepidopteran pheromone blends. We successfully generated a first generation of transgenic plants (SxPv1.0) producing mainly Z11-16OH. Homozygous SxPv1.0 lines maintained the pheromone production up to the T_3_ generation. It should also be noted that basal levels of Z11-16Ald and Z11-16OAc were detected in SxPv1.0, probably produced by endogenous enzymes, since no oxidase was included in this first version of the pathway, and the third enzyme of the route, *EaDAct*, was truncated. The difficulties we experienced in obtaining stable plants producing Z11-16OAc probably indicate that only certain levels/ratios are compatible with viable plant regeneration. The relative position of selection markers in our second-generation construct could have contributed to the recovery of the single plant producing a blend of Z11-16OH and Z11-16OAc. Pheromone production in this new single line (now in the T_2_ generation) is also very stable and maintains remarkably homogeneous levels of production. Having established SxPv1.0 and SxPv1.2 stable plants has allowed us to study in detail the production levels of the different pheromone components, their relative abundance, and their volatility, together with an in-depth characterization of the accompanying phenotype.

As results of our analysis, two main bottlenecks were identified: the associated growth penalty and the poor release rates of the pheromones to the environment. As highlighted also by Reynolds et al., (2017) and later by Xia et al., (2020), one of the most significant downsides to the constitutive overexpression of medium-chain fatty acid biosynthesis pathways in plants is the associated developmental abnormalities. These may result from an imbalance caused by diverting metabolic resources from fatty acid metabolism towards the products of interest, and possibly from the toxicity of the end-products. Such toxic effects can hamper plant viability and result in a negative selection pressure against the genotypes with higher pheromone production levels. Interestingly, whereas Xia et al., (2020) found strong deleterious effects associated to the production of *(E)*-11-tetradecenoic acid, the same authors regenerated normal plants that accumulated Z11-16CoA, the direct precursor of the volatile pheromones produced here. The fact that the simple addition of a desaturase activity leads to deleterious effects may indicate that Z11-16OH itself is responsible for the toxic effects observed when accumulated in leaves. Furthermore, this toxicity seems partially alleviated when a fraction of Z11-16OH is converted to Z11-16OAc, leading to higher biomass in the case of SxPv1.2.

Understanding the changes imposed on the leaf volatilome can shed light on the associated phenotypic changes and the possible imbalances produced by the introduction of the recombinant pheromone pathway. We show here that each SxP version has a distinctive volatile profile, that differs from wild type plants primarily by the presence of the pheromones themselves and a few related fatty-acid derived compounds, which apparently result from endogenous enzyme activities operating on new-to-plant molecules. This seems to be the case for Z11-16Ald (itself a usual component of moth pheromone blends), but also for the differential accumulation of other shorter chain fatty acid derivatives such as 1-pentadecene or hexenal. A close look to the clustered analysis shows that wild type adult *N. benthamiana* tends to produce more monoterpenes (e.g. linalool) and phenolic VOCs (e.g. phenylalanine derivatives) than younger plants. However, this tendency is reduced in SxP plants in general, this effect being even more severe in SxPv1.0 plants. The observed downregulation of the normal volatile components in adult plants could reflect a reduced ability to set up defence mechanisms. Taking place in a generally immune-suppressed species as *N. benthamiana* (Goodin et al., 2008; Bally et al., 2015), this could explain the premature senescence and the early collapse observed in many SxPv1.0 soon after flowering. A strategy to alleviate deleterious effects would consist in disconnecting plant growth from pheromone production. This could be done by employing agronomically-compatible inducible expression systems for the activation of the pathway, taking advantage of the increasingly number of Synthetic Biology tools made available for plants and particularly for *Nicotiana* species (Bernabé-Orts et al., 2020; Cai et al., 2020; Molina-Hidalgo et al., 2020). Alternatively, the use of a different plant chassis displaying specialized structures such as glandular trichomes to store potentially toxic pheromone compounds could be advantageous. Glandular trichomes serve as natural biofactories for VOCs biosynthesis and release, e.g. in aromatic plant species (Huchelmann et al., 2017).

The quantification of pheromone tissue accumulation and environmental release in SxPv1.2 leads to interesting considerations. The maximum pheromone accumulation levels measured in SxPv1.2 reached 174.5 µg g^-1^ FW (totalling both alcohol and acetate forms). This is about half of the levels of precursors reported by Ding et al., (2014) in transient experiments (381 µg g^-1^) or by Xia et al., (2020) in stable plants (335 µg g^-1^), and may indicate a partial conversion into biologically active forms, or upper limit for toxicity, especially in the case of Z11-16OH. However, only a small portion of the plant pheromone content can be detected in the environment after a 72h incubation. Typically, mating disruption strategies require daily release rates between 20 and 500 mg Ha^-1^ day^-1^ (Alfaro et al., 2009; Gavara et al., 2020). Our data indicates that the maximum release rates per biomass unit are around 20 µg Kg plant^-1^ day^-1^, therefore it would require between 1,000 Kg and 25,000 Kg of pheromone-producing intercropping biomass per Ha for effective mate disruption. This is obviously not viable for dwarf SxPv1.2 plants, whose average weight is 9.35 g (aerial parts), and it would be still challenging even if large biomass plant species are used as bio-emitters. Therefore, it is concluded that the improvement in the release rates is an important objective to focus on. In leaves, VOCs are synthesized in mesophyll cells, and release takes place through the stomata or cuticle (Loreto and Schnitzler, 2010). Emission rates of endogenous VOCs are highly variable and depend on the chemical properties of each molecule. Furthermore, volatility is temperature-dependent, with higher temperatures leading to a more rapid transition from the liquid to the gas phase (see Mofikoya et al., (2019) and references herein). C16 fatty acid derivatives are indeed semi-volatile compounds, and in the absence of specialized structures (like the glandular trichomes described above) active transport may play an important role in their release from mesophyll cells. Active transporters of the adenosine triphosphate-binding cassette (ABC) class are known to be required for the release of at least some volatile components of flower scent in petunia (Adebesin et al., 2017). Although molecular mechanisms assisting volatilization of sex pheromone in moth specialized glands it is not well understood, pheromone binding proteins (PBPs) could play an important role in transporting pheromones across the cell membranes into the environment, as well as bringing them in contact with the receptor complexes in the male antennas (Zhou, 2010). The introduction of PBPs or other transporters to facilitate pheromone release needs further exploration. The availability of a first version of a life pheromone bio-dispenser will facilitate the study of transporters and permeability intermediaries and serves as the basis for new design-build-test iterations towards the deployment of efficient SxPs as new components of integrated pest management strategies.

## Materials and Methods

### DNA assembly and cloning

The basic DNA elements (promoters, coding regions, terminators) employed for the assembly of multigene constructs (Level 0 parts) were designed, synthesised and cloned using the GoldenBraid (GB) domestication strategy described in Sarrion-Perdigones et al., (2013). Once cloned into a pUPD2 vector, these new DNA elements were verified by enzymatic digestion and by sequencing. Transcriptional units (Level 1 parts) were then assembled via multipartite BsaI restriction-ligation reactions from level 0 parts, while level >1 modules were produced via binary BsaI or BsmBI restriction–ligation. All level ≥ 1 parts were confirmed by restriction enzyme analysis. All GB constructs created and/or employed in this study are reported in Supplementary Table 1 and their sequences are publicly accessible at https://gbcloning.upv.es/search/features. All constructs were cloned using the Escherichia coli TOP 10 strain. Transformation was performed using the Mix & Go kit (Zymo Research) following the manufacturer’s instructions. The final expression vectors were transformed into electrocompetent Agrobacterium tumefaciens GV3101 C58 or LBA4404 for transient or stable transformations, respectively.

### Transient expression assays in *Nicotiana benthamiana*

*Agrobacterium tumefaciens* GV3101 cultures harbouring the constructs of interest were grown from glycerol stocks for 2 days to saturation, then refreshed by diluting them 1:1000 in LB liquid medium supplemented with the appropriate antibiotics. After being grown overnight, cells were pelleted and resuspended in agroinfiltration buffer (10 mM MES, pH 5.6, 10 mM MgCl_2_ and 200 μM acetosyringone), incubated for 2 hours in the dark, and adjusted to an OD_600_ of 0.1. Equal volumes of each culture were mixed when needed for co-infiltration. A P19 silencing suppressor was included in the mixes to reduce post-transcriptional gene silencing. Agroinfiltration was carried out with a 1 mL needleless syringe through the abaxial surface of the three youngest fully expanded leaves of 4-5 weeks-old plants grown at 24 °C (light)/20 °C (darkness) with a 16:8 h light:darkness photoperiod. Samples were collected 5 days post-infiltration using a Ø 1.5-2 cm corkborer and snap frozen in liquid nitrogen.

### *Nicotiana benthamiana* stable transformation

Stable transgenic lines were generated following the transformation protocol of Clemente (2006), using *Agrobacterium tumefaciens* LBA4404 cultures with the corresponding plasmids. Briefly, leaves from 4-5 -week-old *Nicotiana benthamiana* plants grown at 24 °C (light)/20 °C (darkness) with a 16:8 h light:darkness photoperiod were sterilized by washing in a 2.5% sodium hypochlorite solution for 15 minutes, then rinsed in 70% ethanol for 10 seconds and washed 3 times in sterile distilled water for 15 minutes. Leaf discs were then cut using a Ø 0.8-1.2cm corkborer and transferred to a co-culture medium (MS medium supplemented with vitamins, enriched with 1 mg L^-1^ 6-benzylaminopurine and 0.1 mg L^-1^ naphthalene acetic acid). After 24h on this medium, discs were incubated for 15 minutes in an *Agrobacterium* culture grown overnight to OD_600_ of 0.2 in TY medium (10 g L^-1^ tryptone, 5 g L^-1^ yeast extract and 10 g L^-1^ NaCl, pH=5.6) supplemented with 2mM MgSO_4_·7H_2_O, 200µM acetosyringone and the appropriate antibiotics. After incubation, discs were transferred back to the co-culture medium and incubated for 48h in the dark. Shoots were then induced by transferring to a MS medium supplemented with vitamins, 1 mg L^-1^ 6-benzylaminopurine, 0.1 mg L^-1^ naphthalene acetic acid and 100 mg L^-1^ kanamycin for selection of transformants. After 2-3 weeks of growth with weekly transfers to fresh media, shoots developing from the calli were isolated and transferred to root-inducing medium (MS supplemented with vitamins and 100 mg L^-1^ kanamycin). All in vitro growth was performed in a growth chamber (16:8 h light:darkness photoperiod, 24°C, 60%–70% humidity, 250 lmolm^-2^ s^-1^). Rooted shoots were finally transferred to soil and grown in a greenhouse at 24:20°C (light:darkness) with a 16:8 h light:darkness photoperiod.

### Plant growth and sampling

Transgenic SxP seeds were placed in a germination medium (MS with vitamins 4.9g L^-1^, sucrose 30g L^-1^, Phytoagar 9g L^-1^, pH=5.7) supplemented with 100 mg L^-1^ kanamycin for transgene positive selection. Control WT plants were obtained similarly by placing seeds in a non-selective germination medium. WT and kanamycin-resistant seedlings were transferred to the greenhouse a week after germination, where they were grown at 24:20°C (light:darkness) with a 16:8 h light:darkness photoperiod.

Samples for targeted VOC analysis were collected from the 2^nd^ and 3^rd^ youngest and fully expanded leaves of each plant at the early flowering stage. All samples were frozen in liquid nitrogen immediately after collection, and ground afterwards. Plant size was also estimated at this stage using a 1-10 scale. WT plants grown in parallel with each batch of transgenic plants were taken as reference and given a score of 10.

For the comparative study of the SxPv1.0 and SxPv1.2 lines, seeds from SxPv1.0 5_1_7_X (T_2_), SxPv1.2 4_X (T_0_) and WT *N. benthamiana* plants were all sown simultaneously on selective and non-selective MS medium, then transferred to soil and grown in the conditions described above. Leaf samples and pictures were taken at 4 weeks and 7 weeks after transplant, which corresponds to the young and early flowering stages (herefrom, adults), respectively. Roots were collected at the adult stage. All samples were snap-frozen in liquid nitrogen and ground. All samples were analyzed according to the same GC/MS protocol.

### VOC analysis

50 mg of frozen, ground leaf samples were weighed in a 10mL headspace screw-cap vial and stabilized by adding 1 mL of 5M CaCl_2_ and 150µL of 500mM EDTA (pH=7.5), after which they were sonicated for 5 minutes. Volatile compounds were captured by means of headspace solid phase microextraction (HS-SPME) with a 65 µm polydimethylsiloxane/divinylbenzene (PDMS/DVB) SPME fiber (Supelco, Bellefonte, PA, USA). Volatile extraction was performed automatically by means of a CombiPAL autosampler (CTC Analytics). Vials were first incubated at 80°C for 3 minutes with 500 rpm agitation. The fiber was then exposed to the headspace of the vial for 20 min under the same conditions of temperature and agitation. Desorption was performed at 250 °C for 1 minute (splitless mode) in the injection port of a 6890N gas chromatograph (Agilent Technologies). After desorption, the fiber was cleaned in a SPME fiber conditioning station (CTC Analytics) at 250°C for 5 min under a helium flow. Chromatography was performed on a DB5ms (60 m, 0.25 mm, 1 µm) capillary column (J&W) with helium as carrier gas at a constant flow of 1.2 mL min^-1^. For a first identification of the pheromone peaks, oven programming conditions were 40°C for 2 min, 5°C min^-1^ ramp until reaching 280°C, and a final hold at 280°C for 5 min. Once the target peaks were identified, the oven conditions were changed to an initial 160°C for 2 min, 7°C min^-1^ ramp until 280°C, and a final hold at 280°C for 6 minutes to reduce the overall running time without losing resolution of the desired compounds. Identification of compounds was performed by the comparison of both retention time and mass spectrum with pure standards (for pheromones) or by comparison between the mass spectrum for each compound with those of the NIST 2017 Mass Spectral library (see Supplementary File 1). All pheromone values were divided by the Total Ion Count (TIC) of the corresponding sample for normalization.

The quantification of pheromone compounds emitted by plants was carried out by volatile collection in dynamic conditions. Individual plants were placed inside 5 L glass reactors (25 cm high × 17.5 cm diameter flask) with a 10 cm open mouth and a ground glass flange to fit the cover with a clamp. The cover had a 29/32 neck on top to fit the head of a gas washing bottle and to connect a glass Pasteur pipette downstream to trap effluents in 400 mg of Porapak-Q (Supelco Inc., Torrance, CA, USA) adsorbent. Samples were collected continuously for 72 h by using an ultrapurified-air stream, provided by an air compressor (Jun-air Intl. A/S, Norresundby, Denmark) coupled with an AZ 2020 air purifier system (Claind Srl, Lenno, Italy) to provide ultrapure air (amount of total hydrocarbons <0.1 ppm). In front of each glass reactor, an ELL-FLOW digital flowmeter (Bronkhorst High-Tech BV, Ruurlo, The Netherlands) was fitted to provide an air push flow of 150 mL min^-1^ during sampling. Trapped volatiles were then extracted with 5 mL pentane (Chromasolv, Sigma-Aldrich, Madrid, Spain), and extracts were concentrated to 200 μL under a nitrogen stream. Twenty microliters of an internal standard solution (TFN) were added to the resulting extract prior to the chromatographic analysis for pheromone quantification.

### Statistical analysis

For the untargeted volatilome analysis, data pre-processing was performed with Metalign (Lommen, 2009). Peak intensities were calculated for each compound for the SxP and WT samples and for blanks (mock CaCl_2_ + EDTA samples), and compounds were included in the analysis if the sample:blank ratio was ≥ 2 for at least one of the categories (SxPv1.0, SxPv1.2 or WT). Principal Component Analysis and hierarchical clustering were performed with MetaboAnalyst 5.0 (https://www.metaboanalyst.ca/). After generalized logarithm transformation, data scaling was performed by mean-centering and dividing by the square root of the standard deviation each variable. Hierarchical clustering was done using Ward clustering algorithm and Euclidean distance measure. Plant size values were analyzed with the non-parametric Kruskal-Wallis test using the Past3 software to determine the significance of plant size differences.

### Plant solvent extraction

The total quantity of pheromone compounds accumulated in each plant was extracted with toluene (TLN). Plant samples (ca. 3 g) mixed with fine washed sand (1:1, plant:sand, w/w) were manually ground with a mortar to aid in tissue breakdown and facilitate the extraction. The resulting material was then transferred to 50 mL centrifuge tubes with 10 mL TLN. The extraction process was assisted by magnetic agitation for 12 h and finally by ultrasounds in a Sonorex ultrasonic bath (Bandelin electronic, Berlin, Germany) for 30 min. A 1-mL sample of the resulting extract was taken and filtered through a PTFE syringe filter (0.25 µm). Two-hundred microliters of an internal standard solution were added to the sample prior to the chromatographic analysis for pheromone quantification.

### Synthetic pheromone samples and internal standard synthesis

A synthetic sample of 1g of Z11-16OH was obtained following the method described by Zarbin et al (2007) the sample was carefully purified by column cromatography using silica gel and a mixture of hexane:Et_2_O (9:1 to 8:2) as eluent. Evaporation of the solvent of the corresponding fraction afforded a sample of 96 % purity by GC-FID. Despite its spectroscopical data were fully consistent with those reported by Zarbin et al (2007), some ^13^ C signal described were wrongly assigned or duplicated in the original paper, so we list here a corrected version. Spectroscopical data for Z11-16OH: ^1^H NMR (400 MHz, CDCl3) δ: 0.89 (t, J=6.8 Hz, 3H); 1.22–1.35 (m, 18H); 1.48– 1.60 (m, 2H); 1.95–2.05 (m, 4H); 3.62 (t, J=6.8, 2H); 5.30–5.38 (m, 2H). ^13^C NMR (100 MHz, CDCl3) δ: 13.96, 22.31, 25.70, 26.88, 27.15, 29.26, 29.40, 29.49, 29.52, 29.57, 29.73, 31.93, 32.73, 63.03, 129.85 (2C). MS (70 eV, m/z): 222 (M^+^, 3%), 152 (1%), 137 (4%), 123 (9%), 109 (18%), 96 (48%), 82 (63%), 67 (54%), 55 (100%).

A standard acetylation of Z11-16OH was carried out using acetic anhydride (1.2 eq) and trimethylamine (1.3 eq) as a base in dichloromethane (DCM), affording the corresponding acetate in 95 % yield, whose spectroscopical data where fully coincident with those described in the literature (Zarbin et al. 2007). Oxidation with pyridinium chlorocromate of 100 mg sample of Z11-16OH was carried out following the method described in Zakrzewski et al (2007) affording 62 mg (60 %) of Z11-16Ald, whose spectroscopical data where fully coincident with those described in the literature Zakrzewski et al (2007).

Due to the abundance of compounds structurally related to the pheromone in the biological samples, a straight chain fluorinated hydrocarbon ester (heptyl 4,4,5,5,6,6,7,7,8,8,9,9,9-tridecafluorononanoate; TFN) was selected as the internal standard to improve both sensitivity and selectivity for MS/MS method optimization. TFN was synthesized as follows: to a solution of 4,4,5,5,6,6,7,7,8,8,9,9,9-tridecafluorononanoic acid (500 mg, 1.3 mmol) in DCM, oxalyl chloride was added. After 60 min of continuous stirring, the solvent was removed under vacuum. The residue was re-dissolved in dry DCM (15 ml) and 1-heptanol (0.26 mL, 1.5 mmol) followed by triethyl amine (0.31 ml, 3 mmol) were sequentially added at room temperature, and the resultant solution was refluxed for 24 h. After this period, 15 ml of DCM were added and the solution was successively washed with HCl (1M, 20 ml), NaHCO_3_ (sat., 20 mL), brine (15 ml) and dried with anhydrous MgSO_4_. The solution was filtered and the residue was purified by column chromatography (silica gel; eluent: 1 % Et_2_O/Hexane) to yield heptyl 4,4,5,5,6,6,7,7,8,8,9,9,9-tridecafluorononanoate (281 mg, 45 %), as a colorless oil of 95 % of purity estimated by GC-FID. MS (70 eV, m/z): 393 (10%), 375 (40%), 373 (5%), 132 (10%), 98 (30%), 83 (15%), 70 (100%), 69 (70%), 57 (90%) and 56 (90%).

### Plant extract for biosynthetic pheromone characterization

120 g of SxPv1.2 were mixed with fine washed sand (1:1, plant:sand, w/w) and were manually ground with a mortar to aid in tissue breakdown and facilitate the extraction. The resulting material was then transferred to a 1 L Erlenmeyer flask and 400 mL of hexane were added. The extraction process was assisted by magnetic agitation for 12 h. After this time, the mixture was filtered off and the filtrated was concentrated in a rotary evaporator. The residue (ca. 2 g) was chromatographed in a gravity column (30 cm X 1.5 cm) using silica gel (50 g) as a stationary phase and an mixture Hexane:Et_2_O (9:1) as solvent. 60 fractions of ca. 3 mL were collected and analysed by thin layer chromatography and GC/MS. Those fractions containing biosynthetic Z11-16OH were selected and those containing mainly biosynthetic Z11-16OH where mixed, and the solvent was rotary evaporated, affording 2 mg of material. The ^1^H and ^13^C NMR spectrum of the isolated biosynthetic Z11-16OH was recorded by a Bruker 600 Ultrashield Plus spectrometer (Bruker, Billerica, MA) at a frequency of 600 MHz, using CDCl_3_ as solvent with tetramethylsilane (TMS) as the internal standard.

### Pheromone quantification

The quantification of the pheromone compounds was carried out by gas chromatography coupled to mass spectrometry (GC/MS/MS) using a TSQ 8000 Evo triple quadrupole MS/MS instrument operating in SRM (selected reaction monitoring) mode using electron ionization (EI +), coupled with a Thermo Scientific TRACE 1300 gas chromatograph (GC). The GC was equipped with a ZB-5MS fused silica capillary column (30 m × 0.25 mm i.d. × 0.25 μm; Phenomenex Inc., Torrance, CA). The oven was held at 60 °C for 1 min then was raised by 10 °C min^-1^ up to 110 °C, maintained for 5 min, raised by 10 °C min^-1^ up to 150 °C, maintained for 3 min and finally raised by 10 °C min^-1^ up to 300 °C held for 1 min. The carrier gas was helium at 1 mL min^-1^. For each compound, pheromone components (Z11-16OH and Z11-16OAc) and the internal standard (TFN), the MS/MS method was optimized by selecting the precursor ion and the product ions that provided the highest selective and sensitive determinations (Table 2S).

The amount of pheromone and the corresponding chromatographic areas were connected by fitting a linear regression model, y = a + bx, where y is the ratio between pheromone and TFN areas and x is the amount of pheromone.

### Plant extract fractionation for electroantennography assays

10 g of SxP were mixed with fine washed sand (1:1, plant:sand, w/w) and were manually ground with a mortar to aid in tissue breakdown and facilitate the extraction. The resulting material was then transferred to 50 mL centrifuge tubes with 40 mL TLN. The extraction process was assisted by magnetic agitation for 12 h and finally by ultrasounds in a Sonorex ultrasonic bath (Bandelin electronic, Berlin, Germany) for 60 min. After this time, the mixture was filtered off and the filtrated was concentrated in a rotary evaporator. The residue (ca. 0.2 g) was chromatographed in a gravity column (17 cm X 1 cm) using silica gel (15 g) as a stationary phase and an mixture Hexane:Et_2_O (9:1) as solvent. Twenty-five fractions of ca. 2 ml were collected and analysed by thin layer chromatography and GC-MS. Fractions 17-20 containing biosynthetic Z11-16OH were selected and combined for electroantennography assays.

### Electroantennography assays for evaluating moth response to biosynthetic pheromone

Starter specimens of *Sesamia nonagrioides* (Lefèbvre) (Lepidoptera: Noctuidae) were collected from infested rice (*Oryza sativa* L. (Poales: Poaceae)) plants in paddy fields located in Valencia (Spain). These were maintained in the stems until pupae were obtained and the progeny of the resulting adults was reared on artificial diet (Eizaguirre et al., 1994). Pupae were sexed under stereomicroscope and males were kept separated from females in different chambers under L16:D8 regime at 25±2 °C and 60% relative humidity.

The electrophysiological response of *S. nonagrioides* males to the biosynthetic Z11-16OH was tested by gas chromatography coupled to mass spectrometry and electroantennography detectors (GC/MS-EAD). For this purpose, 2-3 days-old males were individually placed into test tubes in an ice bath to excise their antenna. Between two and five terminal segments of the antenna were also removed with a scalpel. The antenna was mounted between silver wire electrodes impregnated with conductive electrode gel (Spectra 360, Parker Laboratories, Inc., Fairfield, NJ, USA), to increase the electrical contact. A humidified and carbon-filtered airflow (50 ml/min) was directed continuously over the antenna preparation through a glass L-tube placed at less than 2 cm distance. The flow was delivered by a Syntech CS-55 stimulus controller (Ockenfels Syntech GmbH, Kirchzarten, Germany). A pore-sized opening in the elbow part of the L-tube allowed the introduction of the distal part of a fused-silica restrictor connected to the GC apparatus (Clarus 600 GC/MS, Perkin Elmer Inc., Wellesley, PA). The effluent of the GC column (ZB-5MS fused silica capillary column (30 m × 0.25 mm i.d. × 0.25 μm; Phenomenex Inc., Torrance, CA) was split 1:40 for simultaneous detection between the MS and the EAD apparatus. A Swafer S splitter (Perkin Elmer Inc., Wellesley, PA) was employed for this purpose. The GC-MS/EAD run was performed with the SxP extract fraction containing the biosynthetic Z11-16OH obtained as described above. The GC oven temperature was programmed at 120 °C for 2 min, then raised to 200 °C at 10 °C/min and finally from 200 °C to 280 °C (held for 10 min) at 5 °C/min. The EAG responses were recorded with a Syntech IDAC 2 acquisition controller and the GC-EAD 32 (v. 4.3) software was employed for data recording and acquisition (Ockenfels Syntech GmbH, Kirchzarten, Germany).

## Acknowledgments

We want to express our recognition to all the remaining members of the UPV-CSIC Valencia14 iGEM Team, Ivan Llopis, Alejandra González, Alejandro Vignoni, Gabriel Bosque, Maria Siurana, Víctor Nina, Yadira Boada, Alberto Conejero, Javier Urchueguía and Jesús Picó. The iGEM Team received the support of the UPV Generación Espontánea program. Also, thanks to Jaime Primo for his inspiring lectures on the chemistry of insect pheromones. José L. Rambla acknowledges support by a “Juan de la Cierva-Formación” grant (FJCI-2016-28601) from the Spanish Ministry of Economy and Competitiveness. This work was funded by Era-CoBiotech SUSPHIRE and PCI2018-092893 grants.

## BIBLIOGRAPHY

Adebesin, F., Widhalm, J. R., Boachon, B., Lefèvre, F., Pierman, B., Lynch, J. H., et al. (2017). Emission of volatile organic compounds from petunia flowers is facilitated by an ABC transporter. Science 356, 1386–1388. doi:10.1126/science.aan0826.

Agricultural Phermonone Market report FBI100071, 2021 (2021). Global: Fortune Business Insights Available at: https://www.fortunebusinessinsights.com/industry-reports/agricultural-pheromones-market-100071.

Alfaro, C., Navarro-Llopis, V., and Primo, J. (2009). Optimization of pheromone dispenser density for managing the rice striped stem borer, Chilo suppressalis (Walker), by mating disruption. Crop Protection 28, 567–572. doi:10.1016/j.cropro.2009.02.006.

Bally, J., Nakasugi, K., Jia, F., Jung, H., Ho, S. Y. W., Wong, M., et al. (2015). The extremophile Nicotiana benthamiana has traded viral defence for early vigour. Nature Plants 1, 15165. doi:10.1038/nplants.2015.165.

Benelli, Lucchi, Thomson, and Ioriatti (2019). Sex Pheromone Aerosol Devices for Mating Disruption: Challenges for a Brighter Future. Insects 10, 308. doi:10.3390/insects10100308.

Bernabé-Orts, J. M., Quijano-Rubio, A., Vazquez-Vilar, M., Mancheño-Bonillo, J., Moles-Casas, V., Selma, S., et al. (2020). A memory switch for plant synthetic biology based on the phage ϕC31 integration system. Nucleic Acids Research 48, 3379–3394. doi:10.1093/nar/gkaa104.

Bruce, T. J. A., Aradottir, G. I., Smart, L. E., Martin, J. L., Caulfield, J. C., Doherty, A., et al. (2015). The first crop plant genetically engineered to release an insect pheromone for defence. Sci Rep 5, 11183. doi:10.1038/srep11183.

Cai, Y.-M., Kallam, K., Tidd, H., Gendarini, G., Salzman, A., and Patron, N. J. (2020). Rational design of minimal synthetic promoters for plants. Nucleic Acids Research 48, 11845–11856. doi:10.1093/nar/gkaa682.

Carde, R. T., and Minks, A K (1995). Control of Moth Pests by Mating Disruption: Successes and Constraints. Annu. Rev. Entomol. 40, 559–585. doi:10.1146/annurev.en.40.010195.003015.

Cook, S. M., Khan, Z. R., and Pickett, J. A. (2006). The Use of Push-Pull Strategies in Integrated Pest Management. 375–402. doi:10.1146/annurev.ento.52.110405.091407.

Ding, B.-J., Hofvander, P., Wang, H.-L., Durrett, T. P., Stymne, S., and Löfstedt, C. (2014). A plant factory for moth pheromone production. Nat Commun 5, 3353. doi:10.1038/ncomms4353.

Eizaguirre, M., Lopez, C., ASiN, L., and Albajes, R. (1994). Thermoperiodism, Photoperiodism and Sensitive Stage in the Diapause Induction of Sesamia nonagrioides (Lepidoptera: Noctuidae). Journal of Insect Physiology 40(2), 113–119. doi:doi.org/10.1016/0022-1910(94)90082-5.

El-Sayed (2021). The Pherobase: Database of Pheromones and Semiochemicals. Available at: https://www.pherobase.com/.

Evenden, M. (2016). “Chapter 24. Mating Disruption of Moth Pests in Integrated Pest Management: A Mechanistic Approach,” in Pheromone Communication in Moths (University of California Press), 365–394.

Gavara, A., Vacas, S., Navarro, I., Primo, J., and Navarro-Llopis, V. (2020). Airborne Pheromone Quantification in Treated Vineyards with Different Mating Disruption Dispensers against Lobesia botrana. Insects 11, 289. doi:10.3390/insects11050289.

Goodin, M. M., Zaitlin, D., Naidu, R. A., and Lommel, S. A. (2008). Nicotiana benthamiana: Its History and Future as a Model for Plant–Pathogen Interactions. 21, 1015–1026. doi:10.1094/MPMI-21-8-1015.

Hagström, Å. K., Wang, H.-L., Liénard, M. A., Lassance, J.-M., Johansson, T., and Löfstedt, C. (2013). A moth pheromone brewery: production of (Z)-11-hexadecenol by heterologous co-expression of two biosynthetic genes from a noctuid moth in a yeast cell factory. Microb Cell Fact 12, 125. doi:10.1186/1475-2859-12-125.

Holkenbrink, C., Ding, B.-J., Wang, H.-L., Dam, M. I., Petkevicius, K., Kildegaard, K. R., et al. (2020). Production of moth sex pheromones for pest control by yeast fermentation. Metabolic Engineering 62, 312–321. doi:10.1016/j.ymben.2020.10.001.

Hossain, M. S., Williams, D. G., Mansfield, C., Bartelt, R. J., Callinan, L., and Il’ichev, A. L. (2006). An attract-and-kill system to control Carpophilus spp. in Australian stone fruit orchards. Entomologia Experimentalis et Applicata 118, 11–19. doi:10.1111/j.1570-7458.2006.00354.x.

Huchelmann, A., Boutry, M., and Hachez, C. (2017). Plant Glandular Trichomes: Natural Cell Factories of High Biotechnological Interest. Plant Physiol. 175, 6–22. doi:10.1104/pp.17.00727.

Löfstedt, Wahlberg, and Millar (2016). “Chapter 4. Evolutionary Patterns of Pheromone Diversity in Lepidoptera,” in Pheromone Communication in Moths (University of California Press), 43–78.

Lommen, A. (2009). MetAlign: Interface-Driven, Versatile Metabolomics Tool for Hyphenated Full-Scan Mass Spectrometry Data Preprocessing. Anal. Chem. 81, 3079–3086. doi:10.1021/ac900036d.

Loreto, F., and Schnitzler, J.-P. (2010). Abiotic stresses and induced BVOCs. Trends in Plant Science 15, 154–166. doi:10.1016/j.tplants.2009.12.006.

Mofikoya, A. O., Bui, T. N. T., Kivimäenpää, M., Holopainen, J. K., Himanen, S. J., and Blande, J. D. (2019). Foliar behaviour of biogenic semi-volatiles: potential applications in sustainable pest management. Arthropod-Plant Interactions 13, 193–212. doi:10.1007/s11829-019-09676-1.

Molina-Hidalgo, F. J., Vazquez-Vilar, M., D’Andrea, L., Demurtas, O. C., Fraser, P., Giuliano, G., et al. (2020). Engineering Metabolism in Nicotiana Species: A Promising Future. Trends in Biotechnology, S0167779920303073. doi:10.1016/j.tibtech.2020.11.012.

Nešněrová, P., šebek, P., Macek, T., and Svatoš, A. (2004). First semi-synthetic preparation of sex pheromones. Green Chem. 6, 305–307. doi:10.1039/B406814A.

Ortiz, R., Geleta, M., Gustafsson, C., Lager, I., Hofvander, P., Löfstedt, C., et al. (2020). Oil crops for the future. Current Opinion in Plant Biology 56, 181–189. doi:10.1016/j.pbi.2019.12.003.

Petkevicius, K., Löfstedt, C., and Borodina, I. (2020). Insect sex pheromone production in yeasts and plants. Current Opinion in Biotechnology 65, 259–267. doi:10.1016/j.copbio.2020.07.011.

Reynolds, K. B., Taylor, M. C., Cullerne, D. P., Blanchard, C. L., Wood, C. C., Singh, S. P., et al. (2017). A reconfigured Kennedy pathway which promotes efficient accumulation of medium-chain fatty acids in leaf oils. Plant Biotechnol J 15, 1397–1408. doi:10.1111/pbi.12724.

Sans, A., Riba, M., Eizaguirre, M., and Lopez, C. (1997). Electroantennogram, wind tunnel and field responses of male Mediterranean corn borer, Sesamia nonagrioides, to several blends of its sex pheromone components. Entomologia Experimentalis et Applicata 82, 121–127. doi:10.1046/j.1570-7458.1997.00121.x.

Sarrion-Perdigones, A., Vazquez-Vilar, M., Palaci, J., Castelijns, B., Forment, J., Ziarsolo, P., et al. (2013). GoldenBraid 2.0: A Comprehensive DNA Assembly Framework for Plant Synthetic Biology. PLANT PHYSIOLOGY 162, 1618–1631. doi:10.1104/pp.113.217661.

Stelinski, L. L., Gut, L. J., and Miller, J. R. (2013). An Attempt to Increase Efficacy of Moth Mating Disruption by Co-Releasing Pheromones With Kairomones and to Understand Possible Underlying Mechanisms of This Technique. Environ Entomol 42, 158–166. doi:10.1603/EN12257.

Witzgall, P., Kirsch, P., and Cork, A. (2010). Sex Pheromones and Their Impact on Pest Management. J Chem Ecol 36, 80–100. doi:10.1007/s10886-009-9737-y.

Xia, Y.-H., Ding, B.-J., Wang, H.-L., Hofvander, P., Jarl-Sunesson, C., and Löfstedt, C. (2020). Production of moth sex pheromone precursors in Nicotiana spp.: a worthwhile new approach to pest control. J Pest Sci 93, 1333–1346. doi:10.1007/s10340-020-01250-6.

Zakrzewski, J., Grodner, J., Bobbitt, J. M., and Karpińska, M. (2007). Oxidation of unsaturated primary alcohols and ω-haloalkanols with 4-acetylamino-2, 2, 6, 6-tetramethylpiperidine-1-oxoammonium tetrafluoroborate. Synthesis 16, 2491–2494. doi:10.1055/s-2007-983809.

Zarbin, P. H. G., Lorini, L. M., Ambrogi, B. G., Vidal, D. M., and Lima, E. R. (2007). Sex Pheromone of Lonomia obliqua: Daily Rhythm of Production, Identification, and Synthesis. J Chem Ecol 33, 555–565. doi:10.1007/s10886-006-9246-1.

Zhou, J.-J. (2010). “Odorant-Binding Proteins in Insects,” in Vitamins & Hormones (Elsevier), 241–272. doi:10.1016/S0083-6729(10)83010-9.

Zou, Y., and Millar, J. G. (2015). Chemistry of the pheromones of mealybug and scale insects. Nat. Prod. Rep. 32, 1067–1113. doi:10.1039/C4NP00143E.

