## Supplementary Data1 for "Production of Volatile Moth Sex Pheromones in Transgenic *Nicotiana benthamiana* Plants"

**Supplementary data**

**Supplementary Table 1.** GoldenBraid Phytobricks created and used in this study.

| **GB ID** | **Name** | **Description** |
| --- | --- | --- |
| GB1018 | HarFAR CDS | CDS of *Helicoverpa armigera* farnesyl reductase (accession number JF709978) |
| GB1019 | AtrΔ11 CDS | CDS of *Amyelois* *transitella* Δ11-desaturase (accession number JX964774) |
| GB1020 | EaDAct CDS | CDS of the *Euonymus* *alatus* acetyltransferase (accession number GU594061) |
| GB1021 | P35S:HarFAR:T35S | TU for the constitutive expression of *Helicoverpa armigera* farnesyl reductase |
| GB1022 | P35S:EaDAct:T35S | TU for the constitutive expression of *Euonymus alatus* acetyltransferase |
| GB1023 | P35S:AtrΔ11:T35S | TU for the constitutive expression of *Amyelois transitella* Δ11-desaturase |
| GB1024 | P35S:AtrΔ11:T35S-P35S:HarFAR:T35S | Module for the constitutive expression of *Amyelois transitella* Δ11-desaturase and *Helicoverpa armigera* farnesyl reductase |
| GB1025 | P35S:AtrΔ11:T35S-P35S:HarFAR:T35S-SF-P35S:EaDAct:T35S | Module for the constitutive expression of *Amyelois transitella* Δ11-desaturase, *Helicoverpa armigera* farnesyl reductase and *Euonymus alatus* acetyltransferase |
| GB1491 | Tnos:NptII:Pnos-P35S:DsRed:Tnos-P35S:AtrΔ11:T35S-P35S:HarFAR:T35S-P35S:EaDAct:T35S | Module for the constitutive expression of *Amyelois transitella* Δ11-desaturase, *Helicoverpa armigera* farnesyl reductase and *Euonymus alatus* acetyltransferase, together with NptII and DsRed marker genes |
| GB3534 | Omega1_AtrD11+HarFAR+EaDAct+SF | Module for constitutive expression of the Sexy Plant enzymes |
| GB3535 | Omega2_DsRed+nptII | Module for constitutive expression of DsRed and nptII selection genes |
| GB3536 | Alpha1_AtrD11+HarFAR+EaDAct+SF+DsRed+nptII | Module for stable transformation of Sexy Plant genes |
| GB3537 | omega1_DsRed+SF | Construct made to have the DsRed selection gene in an omega plasmid |
| GB3538 | omega2_AtrD11+HarFAR+EaDAct+nptII | Module for constitutive expression of the Sexy Plant genes and the nptII selection marker |
| GB3539 | Alpha1_DsRed+SF+AtrD11+HarFAR+EaDAct+nptII | Module for stable transformation of Sexy Plant genes, flanked by selection markers at both sides |

**Supplementary Table 2.** Optimized values of the MS/MS parameters for each target compound.

| Compound | Transition^1^ | Precursor ion (*m/z*) | Product ion (*m/z*) | Collision energy (eV) |
| --- | --- | --- | --- | --- |
| TFN | 1* | 393 | 375 | 5 |
|  | 2 | 375 | 263 | 10 |
| Z-11-C16:OH  Z-11-C16:OAc | 1 | 82 | 67 | 5 |
|  | 2 | 95 | 67 | 10 |
|  | 3* | 96 | 54 | 10 |
|  | 4 | 96 | 81 | 5 |

^1^ Transitions denoted with (*) were the ones employed to obtain the corresponding chromatographic areas. The others were monitored for confirmatory purposes to have increased selectivity when several peaks appear near to each peak retention time.

**Supplementary Table 3.** Primers created and used in this study for testing the integrity of the EaDAct gene in SxPv1.0 plants.

| **Primer pair name** | **Primer sequences (Fw; Rv)** | **Amplicon fragment size** |
| --- | --- | --- |
| Full | TGCTTCGGCTTCTTTCACTT; GCGATAATGGCAGGGAAGTA | 637 bp |
| CDS | TTGTCTCCCCATAACAATTA; CATGACAACAATATATCACG | 200 bp |
| CHI | CCGGATAGGAATTGGCTAAGATCAT; TGCATTTCACATGCTTGAGTTGACC | 1400 bp |

**(a)	(b)**

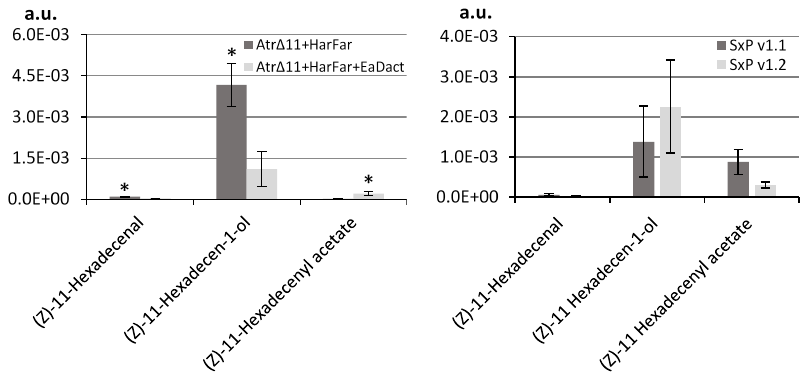

**(c)**
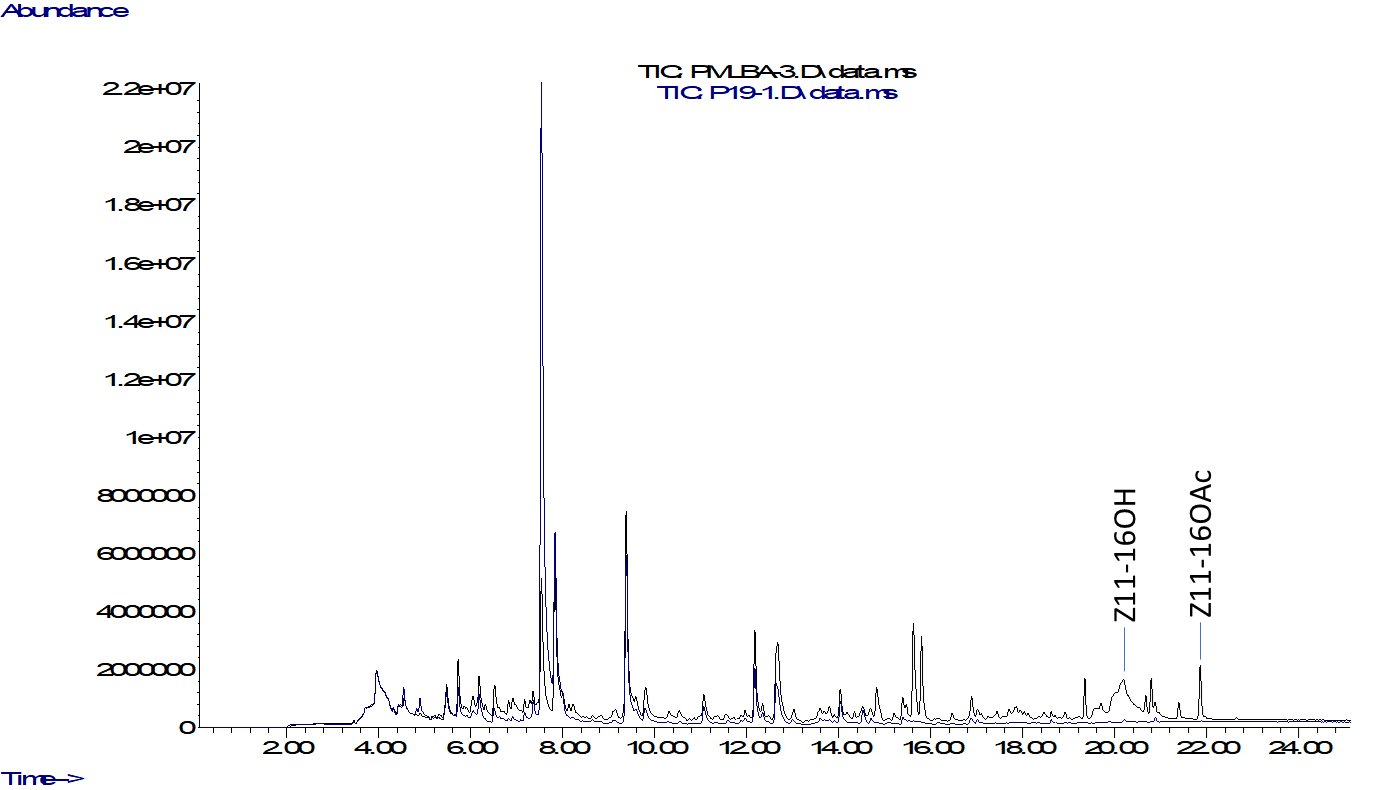

―SP transient

― WT

**Supplementary figure 1.** Transient expression in *Nicotiana benthamiana* of the moth pheromone synthetic pathway. (a) GC/MS quantification of the three pheromones when the three enzymes were transiently expressed, compared to the expression of only the first two enzymes (*AtrΔ11*, *HarFAR*). A 40x increase in acetate levels was observed when EaDAct was present, followed by a 4x decrease in the alcohol levels and a 3x decrease of aldehyde levels (T-test, α=0.05). (b) Pheromone levels obtained transiently with the SxPv1.1 and SxPv1.2 constructs. Error bars represent the ± SE of the measurements of three different leaf samples. (c) A representative chromatogram of a plant agroinfiltrated with the SxPv1.2 construct.

**(a)**

gDNA, Full pair	cDNA, Full pair	_

SP4_2_X	SP7_4_X	SP5_1_X SP5_2_X	WT	C_ C+	SP4_2_X	SP7_4_X	SP5_1_X SP5_2_X	WT	C-	C+

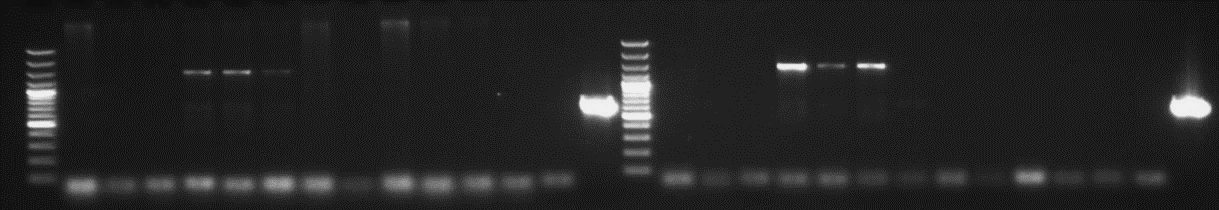

gDNA, CDS pair	cDNA, CDS pair	_

SP4_2_X	SP7_4_X	SP5_1_X SP5_2_X	WT	C-	C+	SP4_2_X	SP7_4_X	SP5_1_X SP5_2_X	WT	C-	C+

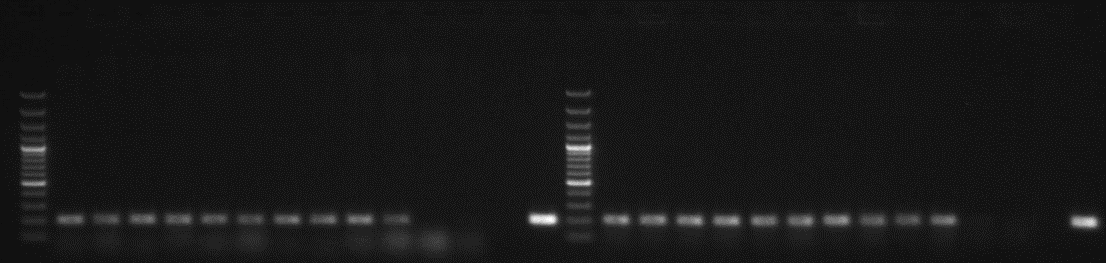

gDNA, CHI pair	cDNA, CHI pair	_

SP4-2-X	SP7-4-X	SP5-1-X SP5-2-X	WT	C-	C+	SP4-2-X	SP7-4-X	SP5-1-X SP5-2-X	WT _	C-	C+

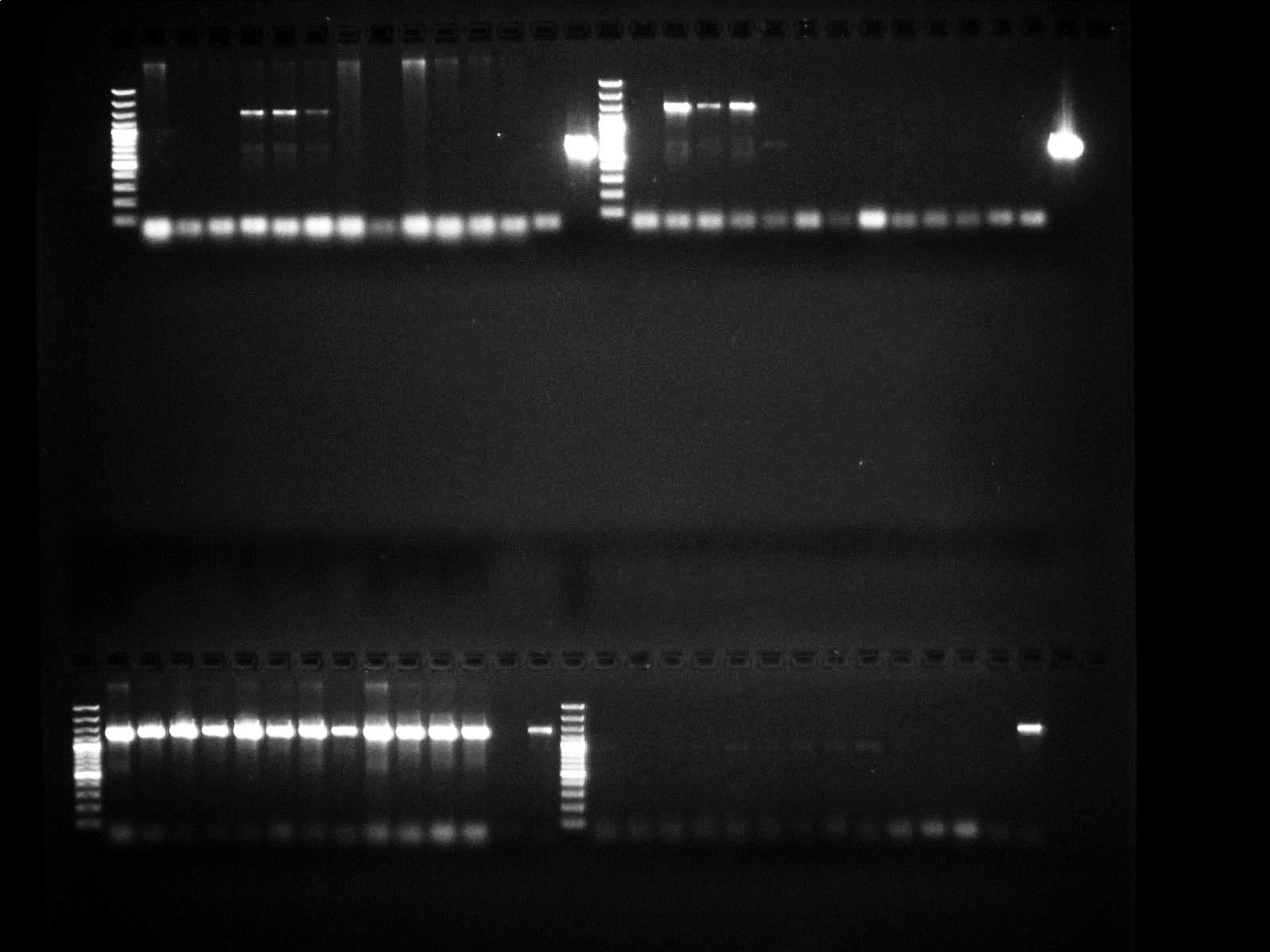

**(b)**

GGACGGTATGGCATTATGATCGCAACTTTGATATCAGCTCATTCATCTTATCAAGTATAACGGGATTTTTTCTAGCTTGGCTTACCACCTTTAAGGTCATTAGCTTTGCATTCGATCAAGGCCCATTATACCCATTACCTCAGAATCTTCTTCATTTTATCTCAATTGCTTGTCTCCCCATAACAATTAAAAGAAATCCAAGCCCAAAATTGAAATCTACAACTAATCCATCACCAATCAGTCATCTTCTTAAAAAAGCCTTTATGAGTTTTCCATCCAAGGTGCTATTCCATTGGGTTATCGCTCATCTGTACCAATACAAAAAATATATGGACCCGAACGTTGTGCTCGTGATATATTGTTGTCATGTTTACGTGGGTGATGCTGCCAACTTACTGATTTAGTGTATGATGGTGTTTTTGAGGTGCTCCAGTGGCTTCTGTTTCTATCAGCTGTCCCTCCTGTTCAGCTACTGACGGGGTGGTGCGTAACGGCAAAAGCACCGCCGGACATCAGCGCTATCTCTGCTCTCACTGCCGTAAAACATGGCAACTGCAGTTCACTTACACCGCTTCTCAACCCGGTACGCACCAGAAAATCATTGATATGGCCATGAATGGCGTTGGATGCCGGGCAACAGCCCGCATTATGGGCGTTGGCCTCAACACGATTTTACGTCACTTAAAAAACTCAGGCCGCAGTCGGTAACCTCGCGCATACAGCCGGGCAGTGACGTCATCGTCTGCGCGGAAATGGACGAACAGTGGGGCTATGTCGGGGCTAAATCGCGCCAGCGCTGGCTGTTTTACGCGTATGACAGTCTCCGGAAGACGGTTGTTGCGCACGTATTCGGTGAACGCACTATGGCGACGCTGGGGCGTCTTATGAGCCTGCTGTCACCCTTTGACGTGGTGATATGGATGACGGATGGCTGGCCGCTGTATGAATCCCGCCTGAAGGGAAAGCTGCACGTAATCAGCAAGCGATATACGCAGCGAATTGAGCGGCATAACCTGAATCTGAGGCAGCACCTGGCACGGCTGGGACGGAAGTCGCTGTCGTTCTCAAATCGTGGGAGCTGCATGACAAAGTCATCGGGCATTATCTGAACATAAAAACACTATCATAGTGGAGTCATTACCCGTTTACGTGATGTGAATATCAGTTGGAGTCTTTGCGCCACTTTAGCAGAGTTCCTTTGTGGTTTTGATGTTGATCCTCAGTTCAAGGAACCATATTTAGCCACTTCTTTGCAGGACTTCTGGGGACGGAGGTGGAACATTATAGTCTCTTCAGTCTTGAGGTCTACTGTTTATGCACCGACGCGTAACATCGCTCAACTCTTCGTCCGTTTTT

**Supplementary figure 2**. Gel electrophoresis of the PCR results with gDNA and cDNA from SxPv1.0 T_2_ plants of the lines SP4_2_X, SP7_4_X and two representative plants for SP5_1_X and SP5_2_X (a) and sequence obtained from the 1.5kb band observed in SP7_4 samples for the Full primer pair (b). Highlighted is the unknown sequence found in the middle of the EaDAct coding sequence. Blast results suggest this sequence could be due to T-DNA re-organizations (data not shown).

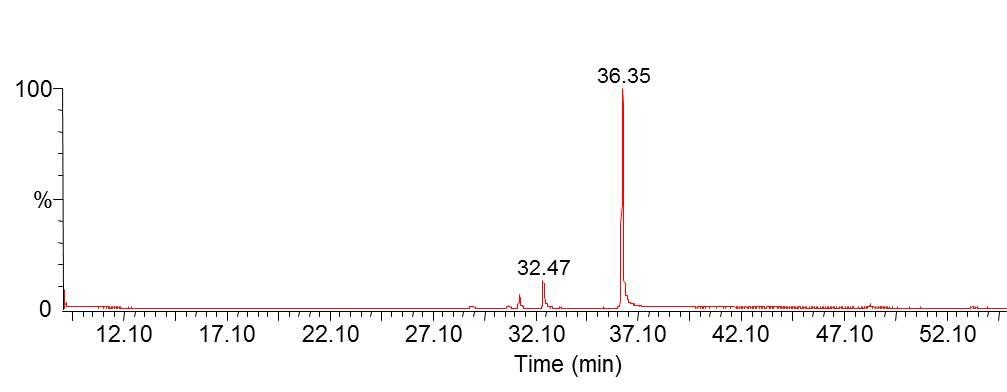

**Supplementary figure 3**. GC/MS of biosynthetic Z11-16OH.

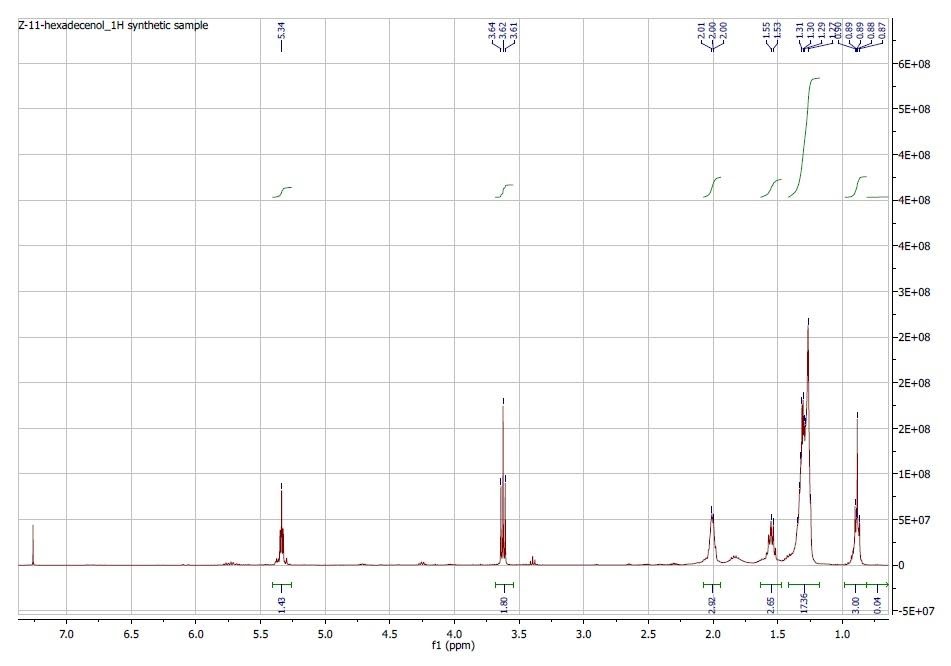

**Supplementary figure 4**. 1H NMR of biosynthetic Z11-16OH.

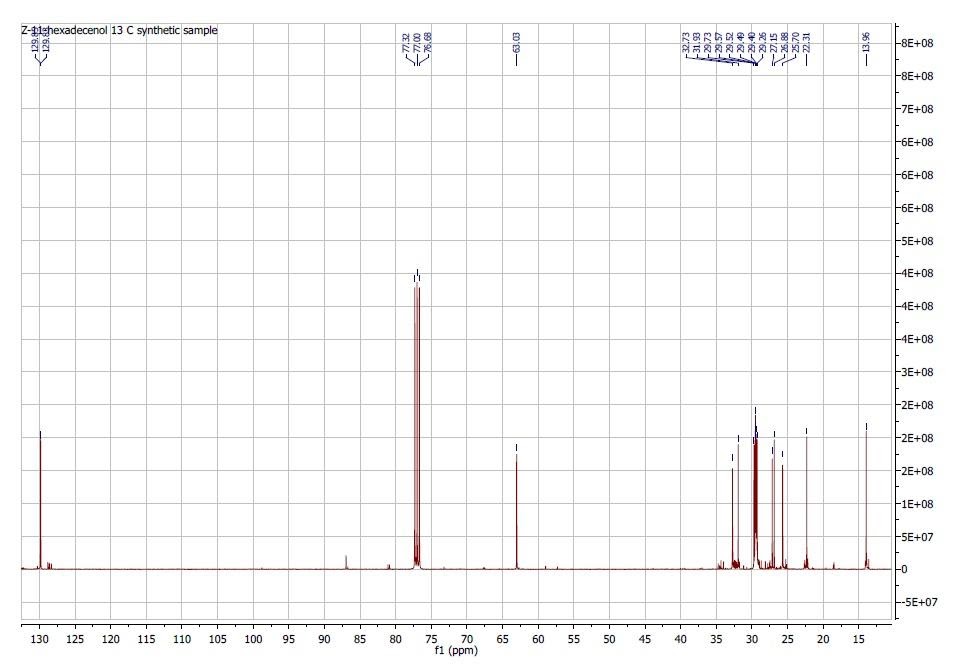

**Supplementary figure 5**. 13C NMR of biosynthetic Z11-16OH.
